# Transcriptomic responses of sponge holobionts to in situ, seasonal anoxia and hypoxia

**DOI:** 10.1101/2023.02.27.530229

**Authors:** Brian W. Strehlow, Astrid Schuster, Warren R. Francis, Lisa Eckford-Soper, Beate Kraft, Rob McAllen, Ronni Nielsen, Susanne Mandrup, Donald E. Canfield

**Author notes:** Denotes co-first author. Author order was decided based on reverse alphabetical order, since the inverse was used on a previous paper. Both authors have the right to list their name first on their CVs.

## Abstract

Deoxygenation can be fatal for many marine animals; however, some sponge species are tolerant of hypoxia and anoxia. Indeed, two sponge species, *Eurypon* sp. 2 and *Hymeraphia stellifera,* survive seasonal anoxia for months at a time. To understand their tolerance mechanisms, we performed differential gene expression analyses on the sponges, their mitochondria and their microbial symbionts under in situ conditions of normoxia, hypoxia and anoxia. Each species possessed a unique microbiome, but the microbiomes of each species were dominated by a species-specific *Thaumarchaeon* and a *Gammaproteobacterium*. Holobiont gene expression was species-and oxygen-level dependent, though there were some shared interspecific responses to deoxygenation. In general, few changes occurred in the expression of sponge metabolic genes as a function of oxygenation level, indicating that they may remain metabolically active under anoxia. However, ATP synthesis genes were significantly upregulated under hypoxia when compared to normoxia, and genes for DNA replication were downregulated. Mitochondrial gene expression was effectively unchanged under both hypoxia and anoxia. Nevertheless, both anoxia and hypoxia caused upregulation of heat shock proteins (HSPs), indicating cellular level adaptations to deoxygenation stress. A meta-analysis demonstrated that sponge transcriptional responses to anoxia were distinct from those displayed by other invertebrates while dormant, and the hypothesis of sponge dormancy under anoxia was not supported. *Thaumarchaeota* symbionts also upregulated stress response genes in hypoxia, while maintaining expression of oxygen-dependent metabolic pathways under hypoxia and anoxia. *Gammaproteobacteria* symbionts showed relatively few noteworthy changes in gene expression in response to anoxia but decreased metabolic gene expression in hypoxia. There was no clear evidence of upregulated anaerobic respiration in the transcriptomes of the sponge holobionts under anoxia or hypoxia. The tolerance of some sponges to prolonged anoxia warrants further investigation and could give them an advantage in future oceans following climate change as well as in ancient oceans when oxygen concentrations were lower than at present.

## Introduction

Ocean oxygen levels are predicted to decline by 1–7% over the next century (Keeling, Körtzinger, and Gruber 2010). Climate change is expanding marine oxygen-minimum zones, while enhanced nutrient runoff is causing coastal eutrophication and subsequent deoxygenation (Kirchman 2021). Deoxygenation is expected to negatively affect the physiology, metabolism and survival of many marine organisms (Diaz and Rosenberg 2008). As a result of deoxygenation and other anthropogenic stressors, we are facing large scale habitat and biodiversity losses, but the full effects of these stressors are still unknown.

Sessile organisms that cannot move or migrate in response to deoxygenation are particularly vulnerable. Sponges (Phylum: Porifera^1^), sessile, ecologically important filter-feeding animals, are hypothesized to suffer from increasing hypoxic/anoxic areas worldwide (Breitburg et al. 2018; Levin 2018; Stramma et al. 2008), and some in situ mortality of sponges (along with other taxa) following deoxygenation has been reported (Altieri et al. 2017; Stachowitsch 1984). However, species-specific responses to deoxygenation in Porifera are variable, and some species are tolerant of hypoxia. Lab experiments have shown that the demosponge *Halichondria (Halichondria) panicea^2^* can survive low oxygen levels down to 0.5–4% of air saturation (a.s.) (Mills et al. 2014), and *Tethya wilhelma* can survive down to 0.25–2% a.s. for 4 days and <0.05% a.s. for at least one hour (Mills et al. 2018). Additional demosponge species that tolerate similar levels of hypoxia include *Geodia barretti^3^* with 7% a.s. (20 µM O2, Leys and Kahn 2018) and *Haliclona pigmentifera^4^* (0.50-0.68% a.s., Gunda and Janapala 2009). In a recent experimental study, four taxonomically distinct sponge species (*Cliona celata*^5^, *Suberites carnosus*^6^, *Polymastia crocea^7^* and *Suberites australiensis*^8^) were exposed to various levels of hypoxia (1.5–40% a.s., 0.13–3.3 mg O_2_ L^-1^) in aquaria. These sponges exhibited species-specific physiological, morphological and behavioral responses to deoxygenation. Sub-lethal oxygen thresholds for these sponges were in the range of 6–20% a.s. (0.48–1.56 mg O_2_ L^-1^) while mortality was only observed in one of the two species (*P. crocea*) exposed to severe hypoxia lower than 5% a.s. (0.13 mg O_2_ L^-1^) (Micaroni et al. 2021a). The sponge species most sensitive to deoxygenation in controlled experiments, *P. crocea*, had a median lethal time (time until death, LT50) of 286 h at ∼1.5% a.s. (0.13 mg O_2_ L^-1^, (Micaroni et al. 2021a), which was still longer than most marine organisms under comparable conditions (reviewed in Micaroni et al. 2021a and Hughes et al. 2020).

Few studies have investigated the deoxygenation tolerance of sponges in situ. Bayer et al. (2020) found that the glass sponge *Vazella pourtalesii^9^* lives in reduced oxygen conditions (< 4 mL/L) on the Scotian Shelf, Canada. Schuster et al. (2021) reported at least nine encrusting demosponge species of the orders Axinellida^10^ and Poecilosclerida^11^ that tolerate seasonal anoxia in a Marine Lake in Lough Hyne, Ireland for ∼three months at a time in the summer, and there are more sponge species at this site (Bell and Barnes 2000). The anoxia-tolerant sponges of Lough Hyne therefore have LT50s that exceed their ca. three month exposure to anoxia, and, compared to other animals, their tolerance is rivaled only by the anemone *Bunodosoma cavernatum^12^* (LT50 in anoxia >1.5 months, Ellington 1982). Moreover, the abundance (percent cover) of encrusting sponges in the seasonally anoxic site of Lough Hyne only slightly decreased over 30 years despite widespread sponge losses in the well oxygenated parts of the lough (Micaroni et al. 2021b), indicating that these sponges are uniquely suited to their seasonally anoxic niche.

Many invertebrates exposed to harsh conditions like anoxia or hypoxia have evolved molecular response pathways to survive in these environments. For instance, under anoxia, marine mollusks shift from lipid, carbohydrate and amino acid metabolism to carbohydrate catabolism as the primary fuel for ATP generation (Larade and Storey 2009). Some mollusks also maximize anoxic survival time (mollusk LT50s at 0 mg L^-1^: 360–672 h, see Micaroni, et al. 2021b) by building up a fixed reserve of internal fuels (primarily glycogen) to increase ATP output and to reduce acidity. A small suite of upregulated genes also plays a critical role in cell preservation, facilitating long-term viability under hypoxia (Larade and Storey 2009). Some anemones may even have elements of a facultative anaerobic metabolism (Sassaman and Mangum 1972).

Only one study has investigated the transcriptomic response of a sponge to hypoxia. After four days of exposure to hypoxia (0.25–2% a.s.), 128 genes within the sponge *T. wilhelma* were differentially expressed compared to controls, but none of these genes were clearly related to metabolic or stress responses (Mills et al. 2018). However, one hour exposure to near anoxia (<0.05% a.s., 105 nM O_2_) elicited stress responses in *T. wilhelma’*s transcriptome (Mills et al. ^2^018^)^, demonstrating that transcriptomic responses were mostly observed under very low oxygen levels.

In most metazoans, the hypoxia-inducible factor (HIF) pathway is essential for maintaining oxygen homeostasis in response to hypoxia (Samenza 2007). However, in sponges and ctenophores, the likely sister-lineages to other metazoans (Feuda et al. 2017), key components of the HIF pathway, including orthologs for HIFa and HIFb, are absent (Mills et al. 2018). HIFa is a transcription factor responsible for triggering metabolic adaptation to deoxygenation in most metazoans (Samenza 2007). Due to the absence of HIFa in sponges, the mechanisms of tolerance to hypoxia and anoxia are largely unknown (Mills et al. 2018).

The sponge microbiome may play a role in sponge deoxygenation tolerance. Sponges can host diverse microbiomes that perform vital functions and can constitute large portions of their biomass (Wilkinson 1978). The host and microbiome together are referred to as a ‘holobiont’ (Webster and Taylor 2012). Some sponge symbionts have anaerobic capabilities, including sulfate reduction (Hoffmann et al. 2005), denitrification, and anaerobic ammonium oxidation (anammox) (Hoffmann et al. 2009). In *V. pourtalesii*, the microbial community is thought to be specialized for the low-oxygen conditions in which it lives (Bayer et al. 2020).

A previous study has shown that the microbiomes of anoxia-tolerant sponges from Lough Hyne were unchanged under anoxia, hypoxia and normoxia despite drastic changes in the microbial communities of the water and sediment, indicating that the holobiont as a whole is tolerant to deoxygenation (Schuster et al. 2021). Specifically, the microbiomes of the anoxia-tolerant sponge species *Eurypon* sp. 2 and *Hymeraphia stellifera^13^* were both dominated by two microbes – one *Thaumarchaeon* and one *Gammaproteobacterium*, though each sponge hosted a different species from each clade. *Eurypon* sp. 2 harbors the *Thaumarchean* OTU1 (closest related isolate: *Nitrosopumilus adriaticus* strain NF5), and *H. stellifera* harbors OTU3 (closest related isolate: *Nitrosopumilus oxyclinae* strain HCE1). For the *Gammaproteobacteria*, *Eurypon* sp. 2 contains OTU2 (closest related isolate: *Arenicella chitinivorans* strain ALEN 1); whereas *H. stellifera* contains OTU7 (closest related isolate: *A. chitinivorans* strain ALEN 1, (Schuster et al. 2021). Although their location within Lough Hyne sponges has not yet been determined, *Gammaproteobacteria* and *Thaumarchaeota* symbionts are abundant and widely distributed throughout the sponge tissue in other species (Moeller et al. 2019). These symbionts also participate in integrated metabolic processes with their host species (Moitinho-Silva et al. 2017).

Schuster et al. (2021) hypothesized potential metabolic pathways within these two symbiont taxa that might function under deoxygenation. Based on the genomes of their close relatives, the *Thaumarchaeota* in these sponges were hypothesized to have components of a facultatively anaerobic metabolism, including fumarate, nitrite, and sulfite respiration and the ability to ferment (Bayer et al. 2020), which could aid in deoxygenation tolerance. A recent study demonstrated that another close *Thaumarchaeota* relative, *Nitrosopumilus maritimus*, can produce trace levels of oxygen (100–200 nM) under environmental anoxia (Kraft et al. 2022).

The pathway of oxygen production and the genes involved are not currently known. However, comparative proteomics suggested that proteins upregulated during oxygen production included some AsnC family proteins and nitroreductases (Kraft et al. 2021). We therefore hypothesized that nitroreductases and AsnC family proteins would be upregulated under hypoxia and anoxia in sponge-associated *Thaumarchaeota. Gammaproteobacteria* symbionts were hypothesized to use sulfide or thiosulfate as an electron donor anaerobically coupled to nitrate reduction, removing exogenous, toxic sulfide from the holobiont under anoxia (Schuster et al. 2021).

To understand the mechanisms of sponge holobiont tolerance and survival under extended anoxia, we performed differential gene expression analysis of the sponges *Eurypon* sp. 2 and *H. stellifera* as well as their respective mitochondria and microbial symbionts including *Thaumarchaeota* and *Gammaproteobacteria*. Sponges were collected from Lough Hyne under in situ normoxia, hypoxia and anoxia. Since neither of these holobionts are established model species, we generated reference transcriptomes for the sponges and assembled the genomes of their symbionts and mitochondria from metagenomic sequences. Additionally, we performed a meta-analysis of the sponge differential expression data to determine if the responses of Lough Hyne sponges to deoxygenation are similar to those of *T. wilhelma* under deoxygenation, other invertebrates under heat stress (Dixon et al. 2015), in a dauer state (Sinha et al. 2012), and during diapause (Gong et al. 2013). Finally, we considered the implications of sponge deoxygenation tolerance for future oceans as well as early animal evolution under low environmental oxygen concentrations.

## Materials and Methods

### Sampling

Sponges were sampled between July 2018 and August 2019 at Labhra Cliff (Lough Hyne Nature Reserve, Ireland; N51°30.05309 W9°18.17679) as described in Schuster et al. (2021) under permit no. R23-27/2018 issued by the Irish Department of Environment, Heritage, and Local Governments. Dissolved oxygen profiles with depth were determined using a Pro20 dissolved oxygen meter (YSI, USA). Profiles for different sampling times are shown in Schuster et al. (2021). Sponge samples were taken using SCUBA (depth range: 15-29 m) under various in situ oxygen conditions that were categorized as follows: normoxic (5.3–12 mg L^-1^, 49.3–111% a.s.), hypoxic (1.30–3.56 mg L^-1^, 12.1–33.1% a.s.), and anoxic (0.00–0.01 mg L^-1^, 0.00–0.93% a.s., i.e. instrument detection limit). Oxygen concentration at each sampling point was verified using a HOBO dissolved oxygen logger (U26-001; Onset, USA). All samples were frozen in liquid nitrogen within ∼15 minutes of collection and kept in a dry shipper until returned to the laboratory where samples were stored in a −80°C freezer until RNA and DNA extraction.

In total 70 sponge samples were taken under anoxia (n = 6), hypoxia (n = 25), and normoxia (n = 39). Although all these red-orange encrusting sponges were indistinguishable in situ, species identification using molecular barcodes revealed that there were nine species (see Schuster et al. 2021). This level of cryptic diversity is common in sponges, and necessitates care in post-sampling species identification. Two species, *Eurypon* sp. 2 and *H. stellifera,* comprised the majority of the samples and were represented across the oxygen conditions; therefore, these two species were selected for the transcriptomic analysis of sponge, symbionts, and mitochondria in this study. For RNA sequencing (RNAseq), eight samples of *H. stellifera* (anoxic = 2, hypoxic = 5, normoxic = 1) were taken, and sixteen samples of *Eurypon* sp. 2 (anoxic = 2, hypoxic = 7, normoxic = 7) were sequenced (n = 24 in total). However, certain oxygen levels within each species, i.e. anoxia in both species and normoxia in *H. stellifera*, had limited replicates (n < 3). The implications of this limited replication are considered in the analysis methods and discussion. Metadata for individual samples, including oxygen concentration, season, depth, collection date, and individual sample code (e.g. DC##) are included in Supplemental Table 1.

### DNA/RNA extraction

Total RNA was isolated using TRIzol^TM^ Reagent from Invitrogen (ThermoFisher Scientific, Denmark), and remaining DNA in the RNA extracts were removed using the DNase MAX kit (Qiagen), following the manufacturers’ protocols.

Additionally, high molecular weight (HMW) DNA was extracted from one individual of each species using the protocol from Vargas et al. (2021) for metagenomic sequencing. The *Eurypon* sp. 2 sample used for metagenomic sequencing was taken under anoxia (sample DC24), and

RNA was also sequenced from this sample. Due to limited availability of samples and DNA, a sample of *H. stellifera* that was not included in the RNAseq (DC297, sampled under normoxia) was used for metagenome sequencing. DNA and RNA yields were determined using a Qubit^TM^ (Thermo Fisher Scientific) and a NanoDrop^TM^ spectrophotometer (Thermo Fisher Scientific). RNA quality was evaluated on an Agilent TapeStation using an RNA ScreenTape (Agilent Technologies).

### Reference transcriptomes and metatranscriptomes

All RNA samples were sequenced twice with two different library preps - one for polyadenylated (poly A) messenger RNA (mRNA, from eukaryotes) and one for total RNA (to also capture prokaryotic RNA). Library preparation for the eukaryotic (sponge) RNA was done using NEBNext® Ultra™ II Directional RNA Library Prep Kit for Illumina (New England Biolabs, USA). For the prokaryotic RNAseq, the supernatant from the poly A enrichment was treated with NEBNext^®^ rRNA Depletion Kit (Bacteria) and then NEBNext^®^ rRNA Depletion Kit (Human/Mouse/Rat, New England Biolabs, USA) to eliminate ribosomal RNA (rRNA) that causes sequence biases. This combination of poly A exclusion and rRNA depletion eliminated the most rRNA compared to other combinations of methods (or single methods in isolation) in preliminary tests, yielding sequences with only ∼25% rRNA. For the eukaryotic (n = 24) and prokaryotic (n = 24) RNA samples, library preparation was completed using the NEBNext Ultra II Directional RNA library preparation kit (New England Biolabs, Denmark). Libraries pooled in equimolar concentrations were sequenced (5M reads per sample, 2 x 50 bp) at the Center for Functional Genomics and Tissue Plasticity (ATLAS) at the University of Southern Denmark (SDU), using a NovaSeq 6000 (Illumina, USA).

To ensure good coverage for the final reference transcriptomes, two samples from each species were sequenced deeper in additional runs. For *H. stellifera*, the sample for deeper sequencing (DC129, sampled under normoxia) was also present in the sample set above. In contrast, for *Eurypon* sp. 2 another sample (DC128, sampled under normoxia) was used due to the limited availability of sample tissue. As in the sequencing above, two libraries were created and sequenced for each sample - one with eukaryotic RNA and one with prokaryotic RNA. For this deeper sequencing, mRNA in the sponge transcriptomes was purified using the NucleoTrap mRNA kit (Macherey-Nagel, Germany). Prokaryotic RNA was rRNA depleted using the Ribo-zero Magnetic kit (Illumina, USA). An additional purification step was performed for all RNA samples using the standard protocol for the CleanNGS kit (CleanNA, Netherlands) before sequencing preparation using the NEBNext Ultra II Directional RNA library preparation kit (New England Biolabs, Denmark). Samples were diluted in equimolar concentrations, pooled and sequenced (2 x 150 bp) on a HiSeq X platform (Illumina, USA) by DNASense ApS (Aalborg, Denmark).

For the eukaryotic RNA, shallow (2 x 50 bp) sequencing generated 9.20 x 10^8^ and 4.38 x 10^8^ reads for *Eurypon* sp. 2 and *H. stellifera*, respectively, across all sampled sponges, and deeper sequences (2 x 150 bp) generated 2.05 x 10^8^ and 3.45 x 10^8^ reads for each sponge of the two species, respectively. For the prokaryotic RNA, shallow sequencing yielded 1.27 x 10^9^ and 6.68 x 10^8^ reads for *Eurypon* sp. 2 and *H. stellifera*, respectively. The deeper prokaryotic sequencing generated 1.52 x 10^8^ and 1.20 x 10^8^ for *Eurypon* sp. 2 and *H. stellifera*, respectively (totals are shown in Supplemental Table 2).

To obtain reference transcriptomes for each sponge species without prokaryotic symbionts, all eukaryotic RNA sequences were combined into a single file. These sequences were trimmed for quality control using public scripts (Matz 2015; Meyer 2016) and assembled with Trinity v 2.8.5 (Grabherr et al. 2011). This assembly was filtered for contaminants and annotated using egg-NOG mapper (Huerta-Cepas et al. 2019). The final sponge transcriptomes included only genes with metazoan annotations or no annotations. For a detailed description of all filtering and assembly steps, see Strehlow et al. (2020) and (https://zenodo.org/badge/latestdoi/573040118) for all scripts used in the current paper. Sponge reference transcriptomes fit within the expected range of size, gene number, and N50 (Supplemental Table 2) of published sponge transcriptomes (Strehlow et al. 2020; Riesgo et al. 2014). Additionally, sponge reference transcriptomes were effectively complete based on Benchmarking Universal Single-Copy Orthologs (BUSCO) analysis (Supplemental Table 2, Simão et al. 2015) and were therefore suitable for use for subsequent differential expression analysis (see below). Sponge reference transcriptome assemblies are available through NCBI under BioProject number PRJNA89319.

### Reference Metagenomes

Two reference metagenomes were sequenced for *Eurypon* sp. 2 and *H. stellifera* at DNASense ApS (Aalborg, Denmark). DNA was fragmented to ∼550 bp (M220 Focused-ultrasonicator, Covaris, USA) and used for library preparation following the NEBNext Ultra II DNA library preparation kit. The DNA library was paired-end sequenced (2 x 150 bp) on a NovaSeq platform (Ilumina, USA).

Sequences were assembled using Megahit v.1.2.9 (Li et al. 2016). The final contig fasta files were used to plot GC (%) vs. mean coverage of mapped reads to identify individual bins. For this, we used the custom python scripts spadescontigstocovgc.py (Francis 2020a) as well as the R script contig_gc_coverage.R (Francis 2020b) to extract bins for the *Thaumarchaeota* and *Gammaproteobacteria* symbiont of each sponge species. Final statistics for each genome are shown in Supplemental Table 2. Raw sequences are available from NCBI under BioProject number PRJNA893197. Metagenome assemblies are available on https://github.com/bstrehlow/Sponge-transcriptomic-responses-to-seasonal-anoxia.

### Mitogenomes

Using only the deep sequences of total (‘prokaryotic’) RNA, the mitochondrial genomes of single individuals of *Eurypon* sp. 2 (DC128) and *H. stellifera* (DC129) were assembled following the pipeline in Plese et al. (2019).

For future utilization, single contigs that likely represent the mitochondrial genomes of *Eurypon* sp. 2 and *Hymeraphia stellifera* were retrieved from the reference metagenome in addition to the coding regions used for differential expression below. The assembled contigs for each metagenome were blasted against the mitochondrial genome of DC128 (*Eurypon* sp. 2) using blastn, with an e-value cutoff of 1×10^-50^ for *Eurypon* sp. 2 (DC97) and 1×10^-30^ for *Hymeraphia stellifera* (DC297). In general, there was only a single hit for each protein coding gene, confirming a single contig as the mitochondrial genome for each sample. Based on the blast, these two contigs contained all protein coding genes retrieved from the transcriptome that were the focus of this analysis. These ‘mitogenomes’ can be accessed here: https://zenodo.org/badge/latestdoi/573040118.

### Differential Expression Analysis

Trimmed reads from the shallow sequencing for each individual were mapped back to their respective reference transcriptome or genome using bowtie2 (version 2.3.5.1, Langmead and Salzberg 2012) and mapped reads were counted for each contig using samcount.pl (see Matz 2015). Eukaryotic reads were mapped to the sponge reference transcriptome for each species. Prokaryotic reads were mapped to their respective metagenomic bins - metagenome assembled genomes (MAGs), i.e. *Thaumarchaeota* or *Gammaproteobacteria*, within each sponge species. The mitochondrial reference was broken into its 14 genes and prokaryotic (total RNA) reads from both species were mapped to their matching mitochondrial reference and counted as above.

Analyses were performed to compare the gene expression of each target (sponge species, symbiont species, and mitochondria; n=8) under the three different oxygen conditions (anoxia, hypoxia and normoxia). Differential gene expression analysis was performed with the package DESeq2 v. 1.22.2 (Love et al. 2016) in R (R Core Team 2016) with (RStudio Team 2020). Three pairwise comparisons of oxygen condition (normoxia vs. hypoxia, normoxia vs. anoxia, and anoxia vs. hypoxia) were computed from DESeq2 results. Significantly differentially expressed genes between each of these pairwise comparisons were identified using a false discovery rate (FDR) adjusted p-value threshold of 0.1. The ReportingTools software package (Huntley et al. 2013) was used to create PDF pages displaying 4 plots of significantly differentially expressed genes and their annotations in various oxygen conditions, as well as box plots, p-values, and LFCs (https://zenodo.org/badge/latestdoi/573040118). These files are searchable in case particular genes are of interest to future studies. We note that comparisons to *H. stellifera* under normoxia were made with only one normoxic sample due to sampling restrictions. Despite this sampling limitation, the normoxic data for *H. stellifera* are still shown as a baseline and reference. For each sponge species, log fold changes (LFC, from DESeq2) in all genes were calculated so that positive LFCs were upregulated and negative values were downregulated in hypoxia or anoxia relative to normoxia. For the comparisons between hypoxia and anoxia, positive LFCs were upregulated in anoxia and vice versa.

### KOG analysis

EuKaryotic Orthologous Groups (KOG class) assignments were used to compare large scale trends and potential correlations in differential gene expression between Lough Hyne sponges under varied oxygen conditions and external gene expression data from other species and conditions. Delta-ranks, a measure of each KOG class enrichment with up or downregulated genes, were calculated from LFC data using the KOGMWU package in R (Dixon et al. 2015), and delta-ranks were plotted using the package pheatmap (Kolde 2013). While it is also possible to calculate delta-ranks with signed p-values (see KOGMWU supporting documentation), LFC was used here to include changes between normoxia and other conditions for *H. stellifera*. Since there was only one normoxic replicant for this species, LFC was a more appropriate metric than signed p values. Within each pairwise comparison (i.e. hypoxic, anoxic, and anoxic vs hypoxic), significantly enriched KOGs were also calculated using KOGMWU (FDR-adjusted p < 0.05).

These KOG enrichments were also compared to those calculated from the sponge *Tethya wilhelma* after four days of exposure to hypoxia and one hour of exposure to anoxia (Mills et al. 2018). To test whether KOG enrichment in sponges exposed to variable oxygen conditions was correlated with gene expression in other invertebrate stress responses, delta rank data was included from adult *Acropora millepora* coral exposed to heat stress (31.5 °C for three days, (Dixon et al. 2015). Similarly, to test the similarity of the sponge data to noted decreases in metabolic activity in invertebrates, KOGs enriched in the dauer (dormant) state in *Caenorhabditis elegans* (Sinha et al. 2012) and the diapause state of the midge *Sitodiplosis mosellana* (Gong et al. 2013) were added using delta rank tables calculated by Strader et al. (2016). Correlations between the KOG delta ranks in different datasets were performed using Pearson’s Correlation in KOGMWU. Within the significantly enriched KOGs ‘replication, recombination and repair’ and ‘energy production and conversion,’ genes that were significantly differentially expressed were plotted in heatmaps to further investigate sponge transcriptomic responses to deoxygenation at the gene level in Lough Hyne sponges.

KOG enrichment analysis was performed as above for each of the two major symbionts within the two sponge species; however, Clusters of Orthologous groups (COGs) were used since they are designed for prokaryotes rather than eukaryotic KOGs and are fewer in number.

### Gene ontology (GO) term enrichment analysis

Log fold changes calculated above were used for rank-based gene ontology (GO) analysis with adaptive clustering (GO_MWU, Wright et al. 2015) to determine what GO terms were significantly enriched under various oxygen conditions. These analyses were run individually for each pairwise comparison of oxygen condition in both sponge species. These analyses were not completed for the two symbiont types because GO terms were not available for these microbes.

### Phylogenetic positions of nitroreductase proteins

To determine if nitroreductases in sponge symbionts are related to genes that may be involved in oxygen production in *N. maritimus,* nitroreductase proteins from the *Thaumarchaeota* symbionts of *H. stellifera* and *Eurypon* sp. 2 were extracted using annotations assigned by egg-NOG. Phylogenetic position of nitroreductase proteins from *H. stellifera* and *Eurypon* sp. 2 were investigated by aligning them to other known nitroreductase proteins from Genbank using Mafft v7 (Katoh and Standley 2013). The ModelTest-NG v.0.1.7 function (Edler et al. 2020) implemented within RaxmlGUI 2.0 was used to calculate the evolutionary model. According to AIC, LG+I+G was chosen as the best-fitting model. Maximum Likelihood (ML) phylogeny was then reconstructed using raxmlGUI v.2.0.6 (Edler et al. 2020) with 100 (rapid) bootstraps. The resulting protein tree was rendered by FigTree v.1.4.4 (http://tree.bio.ed.ac.uk/software/figtree/).

## Results

### Sponge differential expression

There were three pairwise comparisons for significantly differentially expressed genes - hypoxia vs. normoxia, anoxia vs. normoxia, and anoxia vs. hypoxia. The number of significantly differentially expressed genes in these comparisons for sponges, their mitochondria and their symbionts are summarized in Table 1. In the sponge *Eurypon* sp. 2, 737, 1,376 and 1,090 genes were significantly differentially expressed under hypoxia (vs. normoxia), anoxia (vs. normoxia), and anoxia vs. hypoxia, respectively (p < 0.1, Table 1). The differentially expressed genes under each oxygen condition ranged between 1.5–2.8% of the genes of the *Eurypon* sp. 2 reference transcriptome, and 41-48% of these genes are annotated, which reflects the overall annotation percentage of 39.7% for the whole transcriptome (Supplemental Table 2). In hypoxia, the majority of these differentially expressed genes were upregulated (68%). Under anoxia, on the other hand, the majority of genes were downregulated (60%). In anoxia vs. hypoxia, 57% of genes were upregulated in anoxia compared to hypoxia in *Eurypon* sp. 2 (Table 1).

**Table 1.** Number of significantly differentially expressed genes under deoxygenation in the sponge holobionts summarized per reference genome or transcriptome. The direction (up or down) within each paired oxygen condition comparison is relative to normoxia or in the case of anoxia vs. hypoxia, hypoxia. The number of annotated genes is shown in parentheses.

| Sponge | Reference | Total genes in Reference | Hypoxia |  | Anoxia |  | Anoxia vs hypoxia |  |
| --- | --- | --- | --- | --- | --- | --- | --- | --- |
|  |  |  | Up | Down | Up | Down | Up | Down |
| *Eurypon* sp. 2 | sponge | 49,911 | 499<br>(239) | 238<br>(112) | 551<br>(228) | 825<br>(340) | 622<br>(273) | 468<br>(205) |
| *H. stellifera* | sponge | 62,494 | 465<br>(125) | 58<br>(9) | 40<br>(11) | 244<br>(87) | 1531<br>(438) | 623<br>(286) |
| *Eurypon* sp. 2 | mitochondria | 14 | 0<br>(0) | 0<br>(0) | 0<br>(0) | 0<br>(0) | 1<br>(1) | 0<br>(0) |
| *H. stellifera* | mitochondria | 14 | 0<br>(0) | 0<br>(0) | 0<br>(0) | 0<br>(0) | 0<br>(0) | 0<br>(0) |
| *Eurypon* sp. 2 | *Thaumarchaeota* | 1,556 | 19<br>(14) | 1<br>(1) | 3<br>(3) | 0<br>(0) | 0<br>(0) | 0<br>(0) |
| *H. stellifera* | *Thaumarchaeota* | 1,551 | 17<br>(12) | 0<br>(0) | 0<br>(0) | 0<br>(0) | 1<br>(0) | 27<br>(20) |
| *Eurypon* sp. 2 | *Gammaproteobacteria* | 2,751 | 3<br>(3) | 29<br>(29) | 0<br>(0) | 0<br>(0) | 0<br>(0) | 0<br>(0) |
| *H. stellifera* | *Gammaproteobacteria* | 3,057 | 0<br>(0) | 0<br>(0) | 0<br>(0) | 0<br>(0) | 20<br>(20) | 416<br>(401) |

In the sponge *H. stellifera*, 523, 284, and 2,154 genes were significantly differentially expressed in hypoxia, anoxia, and in anoxia vs. hypoxia, respectively (p < 0.1, Table 1). Differentially expressed genes in the different oxygen conditions corresponded to 0.42-3.4% of the total number of genes in the *H. stellifera* transcriptome. Approximately 25-37% of the differentially expressed genes were annotated, which, as in the other sponge species, roughly corresponded to the global percentage (37.0%) of genes annotated within the *H. stellifera* reference transcriptome. Under hypoxia in this species, the vast majority of differentially expressed genes (89%) were upregulated. As in *Eurypon* sp. 2, the majority of differentially expressed genes (84%) were downregulated in anoxia in *H. stellifera*. The majority of differentially expressed genes between anoxia and hypoxia were upregulated in anoxia (71%, Table 1).

#### KOG enrichments in sponges

The global gene expression patterns of Lough Hyne sponges in response to deoxygenation were compared across the two species by assessing up and down regulation of KOG classes (broad functional annotations assigned to genes). Within these sponges, KOG expression in response to hypoxia and anoxia (versus normoxia) were largely distinct in both species (Figure 1A). Expression patterns of both species were only similar in one case between datasets in this study. KOG expression in *H. stellifera* under hypoxia versus anoxia significantly positively correlated with that of *Eurypon* sp. 2 under the same conditions (r = 0.43, p < 0.05, Figure 1E).

**Figure 1.**
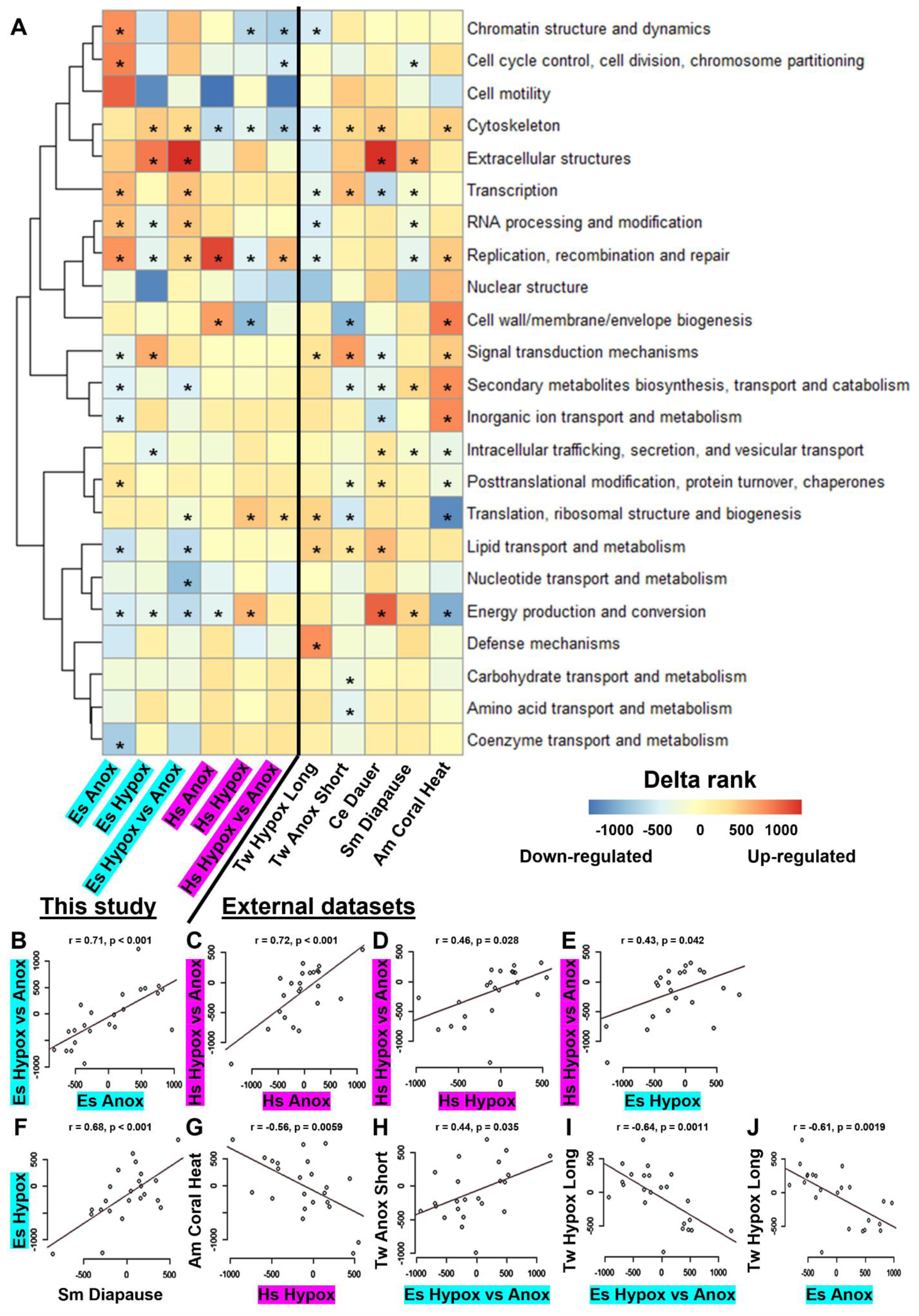
KOG expression profiles of sponges under varied oxygen concentrations compared to each other and external datasets. A. Clustered heatmap (based on correlation) of KOG class (rows) enrichments for upregulated or downregulated genes. Highlighted columns correspond to data generated in this study (Purple: *Hymeraphia stellifera* [Hs], Cyan: *Eurypon* sp. 2 [Es]). Unhighlighted columns represent external datasets. Up or down regulation with respect to normoxia is indicated in ‘Hypox’ (hypoxia) and ‘Anox’ (anoxia) columns, and differential regulation with respect to hypoxia is shown in ‘Hypox vs. Anox’ columns, i.e. upregulated KOGs (red delta ranks) are upregulated in anoxia with respect to hypoxia. Within each column on the heatmap, KOGs that are significantly enriched (FDR-adjusted *p* < .05) are marked with asterisks. KOG enrichments from the sponge *Tethya wilhelma* after four days exposure to hypoxia (Tw Hypox Long) and one hour exposure to anoxia (Tw Anox Short), the coral *Acropora millepora* after three days of exposure to heat stress (Am Coral Heat), *Caenorhabditis elegans* in a dauer state (Ce Dauer), and the midge *Sitodiplosis mosellana* in diapause (Sm Diapause) from external studies were included for comparative purposes. B-E. Significant (*p* < 0.05) correlations between the KOG enrichments for different species and oxygen levels from this study. F-J. Significant correlations between KOG enrichments in this study and those of external datasets. All other possible comparisons between the various datasets were not significant.

Within *Eurypon* sp. 2, KOG class expression patterns in hypoxia vs. anoxia were significantly positively correlated to those of hypoxia (*r* = 0.71, *p* < 0.01, Figure 1B). Similarly, in *H. stellifera*, KOG enrichment in hypoxia vs. anoxia was positively correlated to that of hypoxia (*r* = 0.72, *p* < 0.001, Figure 1C) and anoxia (*r* = 0.46, *p* < 0.05, Figure 1D).

At the level of individual KOGs, the KOG class ‘replication recombination and repair’ was significantly enriched in all pairwise comparisons of oxygen conditions for both sponge species (P_adj_ < 0.05), though in different directions depending on oxygen level. It was significantly upregulated in anoxia and anoxia vs. hypoxia but downregulated in hypoxia in both species (Figure 1A).

In *Eurypon* sp. 2, five genes with replication recombination and repair KOG annotations were significantly upregulated in hypoxia, e.g. HELZ and HELZ2 transcription associated helicases, and 11 genes were significantly downregulated (p < 0.1, Supplemental Figure 4A), including five genes from the minichromosome maintenance (MCM) protein complex. In anoxia, two genes were significantly upregulated within the replication recombination and repair KOG in this species - ‘siz regulation of ARF protein signal transduction’ and DNA polymerase epsilon catalytic subunit (p < 0.1, Supplemental Figure 4B). The majority of significantly differentially expressed replication recombination and repair genes (five) were downregulated in anoxia (p < 0.1, Supplemental Figure 4B). The majority of replication recombination and repair genes (nine) were also downregulated in anoxia vs. hypoxia, with only two significantly upregulated in anoxia (Supplemental Figure 4B). The genes NBN and RFC1 were downregulated in anoxia compared to both normoxia and hypoxia (Supplemental Figure 4B,C). Similarly, the gene ‘siz regulation of ARF protein signal transduction’ was upregulated in anoxia compared to both normoxia and hypoxia (Supplemental Figure 4B,C).

In *H. stellifera*, three genes were significantly differentially expressed in the replication recombination and repair KOG class in hypoxia, and all were upregulated (p < 0.1, Supplemental Figure 4D). No genes within this class were significantly differentially expressed in anoxia (vs. normoxia). Eleven genes were upregulated and eight genes were downregulated in anoxia vs. hypoxia (Supplemental Figure 4E). Similar to *Eurypon* sp. 2, HELZ2 was downregulated in anoxia vs. hypoxia in *H. stellifera*. In contrast to *Eurypon* sp. 2 a distinct MCM gene (MCM6) was downregulated in anoxia vs. hypoxia. Additionally, four genes from the structural maintenance of chromosomes protein 6 (SMC6) complex were downregulated in anoxia vs. hypoxia.

Like replication recombination and repair, the KOG class ‘energy production and conversion’ was significantly enriched (P_adj_ < 0.05) between all conditions except for anoxia vs. hypoxia in *H. stellifera* (P_adj_ = 0.07). The energy production and conversion KOG was significantly upregulated in hypoxia in *H. stellifera* but downregulated in all remaining conditions in both sponge species (Figure 1A).

Within *Eurypon* sp. 2 specifically, four genes were significantly downregulated and twelve genes were upregulated within the energy production and conversion KOG class in hypoxia (p < 0.1, Supplemental Figure 5A). Various nuclear genes for processes in the mitochondria were differentially expressed in hypoxia. SLC25A27 (a mitochondrial membrane uncoupling protein), ndufs8 (a NADH dehydrogenase core subunit in the mitochondrial membrane - complex I), and Surf1 (part of the complex that regulates cytochrome c oxidase assembly) were downregulated in hypoxia (Supplemental Figure 5A). However, other nuclear mitochondrial genes were upregulated, including four subunits of ATP synthase and two additional genes with annotations corresponding to ATP synthesis (Supplemental Figure 5A). In anoxia, fourteen genes were upregulated and five genes were downregulated in the energy production and conversion KOG class (Supplemental Figure 5B).

As in hypoxia, ndufs8 was downregulated, but no other significant differentially expressed genes were shared across the two oxygen levels. The following genes were also downregulated in anoxia: mitochondrial sulfite oxidase (SUOX), malate dehydrogenase (MDH1), and two genes belonging to the ATP gaunido phosphatase family (Supplemental Figure 5B). Conversely, four genes involved in phosphoenolpyruvate carboxykinase activity were significantly upregulated in anoxia (p < 0.1, Supplemental Figure 5B). Additionally, three mitochondrial membrane transporter proteins from the SLC family, two CYB5R4 cytochrome reductase genes, a mitochondrial carrier protein, and fdx1 cytochrome reductase were upregulated in anoxia. In anoxia vs. hypoxia in *Eurypon* sp. 2, ten genes were upregulated in anoxia (Supplemental Figure 5C). Upregulated genes included all genes that were significantly upregulated in anoxia, except for the phosphoenolpyruvate carboxykinase associated genes. Six genes were significantly downregulated in anoxia vs. hypoxia. Of these, three were also downregulated in anoxia (vs. normoxia) and two genes were significantly upregulated in anoxia (Supplemental Figure 5A-C). It is noteworthy, however, that all genes upregulated genes in anoxia and hypoxia were only strongly upregulated in one of the two anoxia replicates (DC56, Supplemental Figure 5 B,C).

In *H. stellifera*, eight genes were upregulated and two genes were downregulated in hypoxia. As in *Eurypon* sp. 2, the ATP5A1 ATP synthase subunit was upregulated in hypoxia. An additional ATP synthase subunit (ATP6V0C) that was not significantly upregulated in *Eurypon* sp. 2 was upregulated in *H. stellifera* in hypoxia (p < 0.1, Supplemental Figure 5D). Mitochondrial sulfide:quinone oxidoreductase (SQRDL), three genes associated with phosphoenolpyruvate carboxykinase, and a histone demethylase (KDM6A) were also significantly upregulated in hypoxia in *H. stellifera*. Conversely, PCK1 (also associated with phosphoenolpyruvate carboxykinase) and MDH1 were significantly downregulated in hypoxia (p < 0.1, Supplemental Figure 5D). In anoxia in *H. stellifera*, ATP5A1 was significantly upregulated, and ATP6V0C, PCK1, and MDH1 were significantly downregulated (Supplemental Figure 5E). Comparing anoxia vs. hypoxia, eight genes were upregulated and five genes were downregulated (Supplemental Figure 5F).

As in hypoxia, SQRDL, four genes associated with phosphoenolpyruvate carboxykinase activity, and KDM6A were upregulated in anoxia vs. hypoxia. Additionally, two unshared genes, including the mitochondrial Succinyl-CoA:3-ketoacid coenzyme A transferase 1 (OXCT1) and a mitochondrial carrier protein, were upregulated in anoxia vs. hypoxia (Supplemental Figure 5F). Five genes were downregulated in anoxia vs. hypoxia in *H. stellifera*. Two of these four only exhibited high expression levels in one hypoxic sample (DC105) that drove the observed difference (Supplemental Figure 5F). The other three downregulated genes in anoxia vs. hypoxia were mitochondrial creatine kinase (CKMT2), NAD P transhydrogenase (NNT), and proto-oncogene tyrosine-protein kinase (Src).

The ‘cytoskeleton’ KOG class was also significantly enriched (P_adj_ < 0.01) in all but one oxygen condition – anoxia in *Eurypon* sp. 2 (P_adj_ = 0.1). Interestingly, enrichment patterns in this KOG were species specific, being significantly upregulated in hypoxia and anoxia vs. hypoxia in *Eurypon* sp. 2 and downregulated in all conditions for *H. stellifera*.

The other twenty KOG classes were only significantly enriched in Lough Hyne sponges under a few (or no) different oxygen conditions and patterns of significant enrichment were generally condition and species-specific (Figure 1A). With the exception of energy production and conversion, KOGs pertaining to metabolic processes were rarely significantly enriched. Expression of the KOG classes ‘carbohydrate metabolism’ and ‘amino acid transport and metabolism’ were not significantly changed under any oxygen condition (P_adj_ > 0.05). There was some significant enrichment of metabolic KOGs, but only in *Eurypon* sp. 2. ‘Coenzyme transport and metabolism’ and ‘inorganic ion transport and metabolism’ were both downregulated in anoxia, and ‘nucleic acid transport metabolism’ was downregulated in hypoxia vs. anoxia. For both these conditions (anoxia and hypoxia vs. anoxia), ‘Lipid transport and metabolism’ and ‘Secondary metabolites biosynthesis, transport and catabolism’ were significantly downregulated. However, the transcription of the majority of genes labeled with metabolic KOGs remained unchanged under anoxia and hypoxia.

Comparing the Lough Hyne dataset with external datasets of invertebrate KOG expression in response to stress, only two positive correlations were found. In *Eurypon* sp. 2, expression patterns under anoxia vs. hypoxia were comparable to those of *T. wilhelma* under short-term anoxia (*r* = 0.44, *p* < 0.05, Figure 1H). The hypoxic response of *Eurypon* sp. 2 resembled that of diapause in *S. mosellana* (*r* = 0.68, *p* < 0.001, Figure 1F), but this was the only correlation found between Lough Hyne sponge data and diapause or dauer states. Importantly, KOG enrichments under anoxia (versus normoxia) were not positively correlated to any other external datasets for either sponge species.

Three significant negative correlations, which corresponded to opposing KOG expression patterns, were observed between Lough Hyne sponges and external datasets. Hypoxia responses in *H. stellifera* were negatively correlated with those of *A. millepora* under heat stress (*r* = -0.56, *p* < 0.01, Figure 1G). The long exposure of *T. wilhelma* to hypoxia was negatively correlated to both anoxia (*r* = -0.64, *p* < 0.01, Figure 1J) and hypoxia vs. anoxia (*r* = -0.61, *p* < 0.01, Figure 1I) in *Eurypon* sp. 2.

#### GO term enrichments in sponges

Generally, significantly enriched GO terms between the different treatments were not shared between the two sponge species (Supplemental Figures 1-3). The strongest upregulated GO enrichments (*p* < 0.01) in hypoxia in *Eurypon* sp. 2 were cellular signaling and ‘lipoprotein particle receptor binding’ (Supplemental Figure 1A). For *H. stellifera*, on the other hand, the strongest upregulated enrichments (*p* < 0.01) in hypoxia included the terms ‘protein tyrosine kinase binding,’ ‘molecular adaptor,’ ‘structural molecule,’ and ‘structural constituent of ribosome’; whereas the downregulation of the term ‘motor’ was strong (*p* < 0.01, Supplemental Figure 1B).

Under anoxia, strong upregulation of the ‘proteasome binding’ term was observed in *Eurypon* sp. 2 (*p* < 0.01), and the terms ‘phosphoenolpyruvate carboxykinase’, ‘carbohydrate kinase,’ monovalent inorganic cation transmembrane transporter,’ and ‘sodium ion transmembrane transporter’ were highly downregulated (*p* < 0.01, Supplemental Figure 2A). In *H. stellifera* under anoxia, the terms ‘damaged DNA binding’ and ‘calcium-dependent phospholipid binding’ were highly upregulated, and four terms associated with various aspects of the cytoskeleton were downregulated (*p* < 0.01, Supplemental Figure 2B).

In *Eurypon* sp. 2 under anoxia vs. hypoxia, the terms ‘lipoprotein receptor binding’ and ‘proteasome binding’ were strongly upregulated in anoxia, but no terms were strongly downregulated in anoxia versus hypoxia (*p* < 0.01, Supplemental Figure 3A). The former term was also upregulated under anoxia vs. normoxia. Conversely, the strongest enrichments in *H. stellifera* in anoxia vs. hypoxia were downregulated in anoxia and were composed of three of the four cytoskeletal binding GO terms that were also downregulated in anoxia vs. normoxia (see above, *p* < 0.01, Supplemental Figure 3B), but no strong upregulation of GO terms was observed.

Considering GO enrichments at lower significance levels (0.01 < *p* < 0.1), within *Eurypon* sp. 2, the GO terms ‘proteasome binding’, ‘lipoprotein particle receptor binding,’ and ‘Hsp70 binding’ (HSP: Heat shock protein) were significantly upregulated in all cases of decreased oxygen albeit at varied significance levels (*p* < 0.05, Supplemental Figures 1–3A). Between only two levels, the terms ‘transmembrane signaling receptor,’ ‘secondary active transmembrane transporter,’ and ‘molecular transducer’ were upregulated in hypoxia but downregulated in anoxia. However, the term lipoprotein particle receptor binding was downregulated in anoxia and upregulated in hypoxia (*p* < 0.1, Supplemental Figures 1,2A). Additionally, the term ‘structural constituent of ribosome’ was upregulated in anoxia but downregulated in anoxia vs. hypoxia (*p* < 0.1, Supplemental Figures 1,3A). No additional GO terms other than the three present in all oxygen levels for *Eurypon* sp. 2 were enriched in both the pairwise comparison of anoxia and anoxia vs. hypoxia.

**Figure 2.**
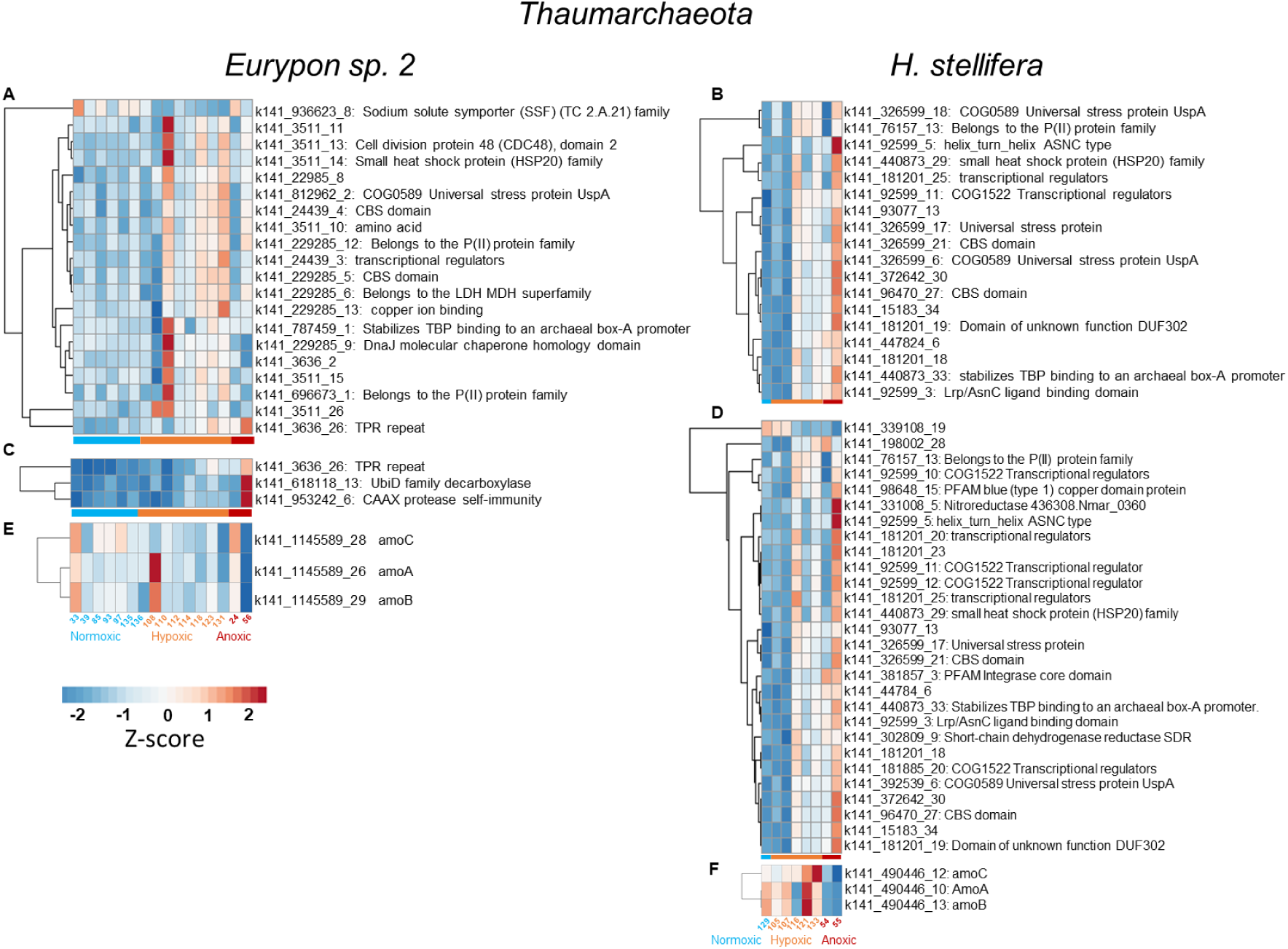
A-D. All significantly differentially expressed genes for the *Thaumarchaeota* in both sponge species (left: *Eurypon* sp. 2, right: *H. stellifera*. For *Thaumarchaeota* associated with *Eurypon* sp. 2, only two pairwise comparisons of oxygen conditions had significant differences: A. hypoxia vs. normoxia and C. anoxia vs. normoxia. In symbionts associated with *H. stellifera*, there were significantly differentially expressed genes in hypoxia vs. normoxia (B) and anoxia vs. hypoxia (D). There were no significant differences in the three genes involved in ammonia oxidation (AmoA, B and C) in either symbiont, but their heatmaps are shown in E and F for *E.* sp 2 and *H. stellifera*, respectively.

**Figure 3.**
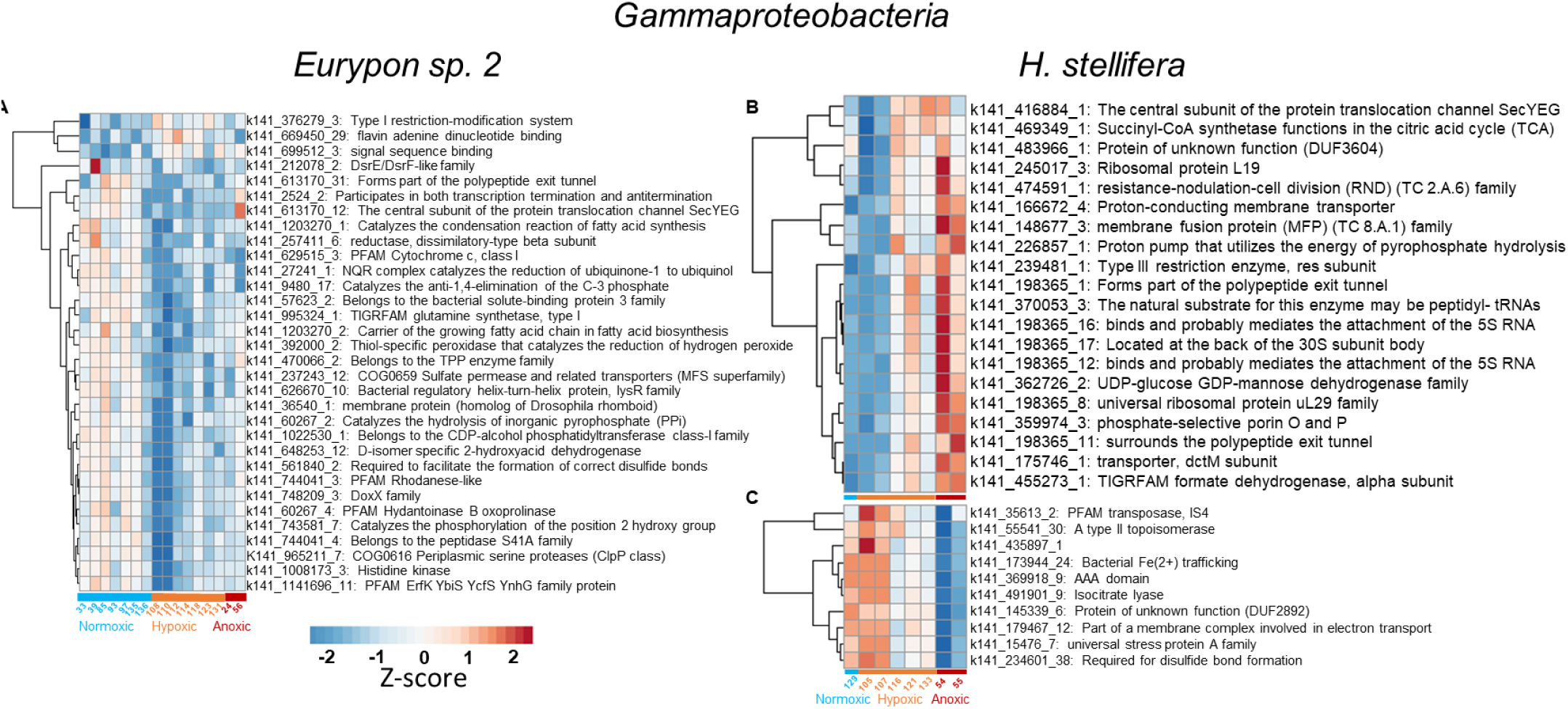
All significantly differentially expressed genes for the *Gammaproteobacteria* in both sponge species (A: *Eurypon* sp 2, B: *H. stellifera*. For *Gammaproteobacteria* associated with *Eurypon* sp. 2, there were only significant differences between hypoxia vs. normoxia (A). In symbionts associated with *H. stellifera*, there were only significantly differentially expressed genes in anoxia vs. hypoxia (B,C). All downregulated genes in anoxia vs. hypoxia are shown in B. There were 402 genes that were downregulated in anoxia vs. hypoxia in *Gammaproteobacteria* from *H. stellifera*, so only the top 10 based on p value are shown (C).

No GO terms were significantly up or downregulated in all three levels of oxygen conditions for *H. stellifera*. However, some GO terms were significantly enriched in two levels. The term ‘ATPase, coupled to transmembrane movement of ions, rotational mechanism’ was downregulated in hypoxia but upregulated in anoxia (*p* < 0.1, Supplemental Figures 1B, 2B respectively). The GO terms ‘I-SMAD binding’, thioesterase binding,’ and ‘ubiquitin-like protein ligase binding’ were all upregulated in hypoxia (vs. normoxia) and anoxia vs. hypoxia, while the ‘motor’ term was downregulated (*p* < 0.1, Supplemental Figures 1B, 3B).

There were only two shared GO enrichments between both sponge species for any given oxygen condition. The terms ‘protein tyrosine kinase binding’ and ‘structural constituent of the ribosome’ were upregulated in both species in hypoxia (*p* < 0.1, Supplemental Figure 1), although they were enriched to different degrees of significance. No significantly enriched GO terms were shared between species in normoxia vs. anoxia or hypoxia vs. anoxia.

At the gene level, significantly differentially expressed heat shock proteins (Hsps) in both sponges are shown in Supplemental Figure 8. In *Eurypon* sp. 2, one Hsp gene (DNAJC8/Hsp40) was downregulated in hypoxia (p < 0.01), but four Hsp genes (HSPA8, two genes for HSP90B1, and Hsp70) were significantly upregulated in hypoxia (p < 0.1, Supplemental Figure 8A). Additionally, HSPA8 and C11orf73 (Hsp70 binding protein) were upregulated in anoxia vs normoxia (p < 0.05, Supplemental Figure 8A). Another Hsp70 gene was also significantly downregulated in anoxia vs. hypoxia (p < 0.05, Supplemental Figure 8A). In *H. stellifera*, HSP90A1 was upregulated in hypoxia and anoxia (p < 0.1, Supplemental Figure 8B). Two additional Hsp genes (both for HSP90B1) were also upregulated in anoxia. Finally, the two HSP90B1 genes were upregulated in anoxia vs. hypoxia, but HSPA5 was downregulated in anoxia vs. hypoxia (p < 0.05, Supplemental Figure 8B).

### Mitochondrial gene expression

All 14 coding genes from the sponge mitochondria were transcribed in both species and in all oxygen conditions. There was only one mitochondrial gene (nad2) that was significantly differentially expressed but only in *Eurypon* sp. 2. This gene was significantly upregulated (p < 0.1) in anoxia vs hypoxia (Supplemental Figure 6A, Table 1). Similar counts for each individual mitochondrial gene across both sponge species and all oxygen conditions were found. For instance, counts for atp6 were consistently twice as high as counts for atp8 (Supplemental Figure 6).

### Prokaryotic differential expression

#### Thaumarchaeota gene expression

For the *Thaumarchaeota* associated with *Eurypon* sp. 2, 20, 3 and 0 genes were differentially expressed in hypoxia, anoxia, and hypoxia vs. anoxia, respectively (p < 0.1, Table 1). These genes accounted for 0-1.3% of this *Thaumarchaeota* genome, and 75-100% of the differentially expressed genes were annotated (Table 1), compared to 69.2% annotated in the entire reference genome. All but one of the differentially expressed genes in hypoxia were upregulated (Table 1, Figure 2A). The downregulated gene belonged to the sodium solute symporter family (SSF, TC 2.A.21, Figure 2A). The upregulated genes in hypoxia included three stress response genes (COG0589 Universal stress protein [UspA], small heat shock protein [HSP20], and DnaJ molecular chaperone homology domain), two CBS domains, a member of the lactate/malate dehydrogenase (LDH MDH) superfamily, a tetratricopeptide (TPR) repeat, two transcription regulators, and a PII signal transduction protein.

Only three genes were significantly differentially expressed in anoxia in *Eurypon* sp. 2’s *Thaumarchaeota*, and all were upregulated (Table 1, Figure 2C). The upregulated genes in anoxia were a UbiD family decarboxylase, CAAX protease self-immunity, and the same TPR repeat that was also upregulated in hypoxia. There were no significantly differentially expressed genes between anoxia and hypoxia in this *Thaumarchaeota* (Table 1). The genes necessary for ammonia oxidation, i.e. subunits A, B and C of ammonia monooxygenase (Amo), were present in this *Thaumarchaeota*, but they were not differentially expressed under deoxygenation (Figure 2E).

In the *Thaumarchaeota* symbiont of *H. stellifera*, 17, 0 and 28 genes were significantly differentially expressed in hypoxia, anoxia and hypoxia vs. anoxia, respectively (Table 1). Similar to the symbiont of *Eurypon* sp. 2, all 17 differentially expressed genes in the *H. stellifera* symbiont were upregulated in hypoxia. These genes included the stress response genes ‘universal stress protein, an HSP20 family protein, and two copies of UspA. HSP20 and UspA were also upregulated in the *Eurypon* sp. 2 holobiont. Similarly, a PII signal transduction protein, CBS domain, and two transcription regulators (one identical: TBP stabilizer, and one distinct to *H. stellifera* symbionts) were upregulated in both *Thaumarchaeota* species in hypoxia. Unlike in *Eurypon* sp. 2, the *Thaumarchaeota* in *H. stellifera* upregulated a helix turn helix ASNC type domain and a Lrp/AsnC ligand binding domain (Figure 2B). No genes were significantly differentially expressed in anoxia in this *Thaumarchaeon*.

As in the other holobiont species, *Thaumarchaeota* genes for ammonia oxidation were not significantly differentially expressed with *H. stellifera* symbionts (Figure 2F). Twenty-seven genes were upregulated and one gene (K141_339108_19, unannotated) was downregulated in anoxia vs. hypoxia in the *Thaumarchaeota* within *H. stellifera* (Figure 2D). Most of the upregulated genes in anoxia vs. hypoxia were also upregulated in hypoxia. Upregulated genes in anoxia vs. hypoxia unique to the *H. stellifera* holobiont included additional transcriptional regulators, an integrase domain, and a putative nitroreductase.The phylogenies of this upregulated nitroreductase and all nitroreductases from the studied holobionts as well as publicly available nitroreductase sequences from other *Thaumarchaeota* are shown in Supplemental Figure 9.

For both *Thaumarchaeota*, only two COGs (‘signal transduction mechanisms’ and ‘transcription’) were significantly enriched (P_adj_ < 0.05, Supplemental Figure 7A). These two COGs were both downregulated in hypoxia and anoxia vs. hypoxia within *H. stellifera* symbionts. All other COG classes were comparable in their expression levels regardless of holobiont species or oxygen concentration (Supplemental Figure 7A).

#### Gammaproteobacteria gene expression

The *Gammaproteobacteria* associated with *Eurypon* sp. 2 only exhibited significant differential expression under hypoxia but not in any other oxygen condition (Table 1). In hypoxia, three genes were upregulated: type 1 restriction-modification system, flavin adenine dinucleotide binding, and signal sequence binding (Figure 3A). The 29 downregulated genes in hypoxia included two genes involved in sulfur processing (k212078_2, k237243_12), cytochrome c (class I), a gene associated with the assembly of the periplasmic c-type cytochromes (k561840_2), a NQR complex gene, various genes involved in pore formation, a Rhodanese-like gene, glutamine synthetase (type I), and YbiS YcfS YnhG family protein.

In the *Gammaproteobacteria* associated with *H. stellifera*, genes were only significantly differentially expressed between anoxia and hypoxia (436 genes, p <0.1, Table 1, Figure 3B,C). Twenty genes were upregulated in anoxia vs. hypoxia and all were annotated (Figure 3B). Upregulated genes in anoxia vs. hypoxia included five genes associated with the ribosome (k141_198365_15, k141_198365_8, k141_198365_17, and k141_245017_3), various genes for molecular transport, a transmembrane proton pump (k141_226857_1), succinyl-CoA synthetase, and formate dehydrogenase (alpha subunit). There were also 416 genes that were significantly downregulated in anoxia vs. hypoxia (Table 1). The top ten upregulated genes (lowest p values) are exhibited in Figure 3C.

*Gammaproteobacterial* COG expression is shown in Supplemental Figure 7B. Only one COG class, ‘translation, ribosomal structure and biogenesis,’ was significantly enriched in *Gammaproteobacteria* under the different oxygen conditions (P_adj_ < 0.05). This COG class was downregulated in *Gammaproteobacteria* associated with *Eurypon* sp. 2 under hypoxia as well as those of *H. stellifera* in anoxia vs. hypoxia (Figure 7B).

## Discussion

Sponge holobiont gene expression was significantly changed by exposure to deoxygenation in situ, though responses were generally species-specific, dependent on oxygen level, and varied across holobiont members. The potential functions of this differential expression are discussed below for each member of the two holobionts: sponge, sponge mitochondria, *Thaumarchaeota* and *Gammaproteobacteria*. Changes in gene expression within each member of the holobiont is discussed in a separate section below, and then the potential functional adaptations to deoxygenation at the level of the holobiont are summarized. The implications of this study for past and future oceans are considered.

### Differential expression in sponges during deoxygenation

Cellular stress responses were observed in the expression of HSPs, a group of stress response genes that have been widely studied in marine invertebrates. These proteins are molecular chaperones involved in protein folding, unfolding, transport, assembly of protein complexes, and the protection of restoration of proteins exposed to stressors, and they are present in nearly all organisms (Li and Srivastava 2004). Both Lough Hyne sponges upregulated HSP genes in response to hypoxia and anoxia (vs normoxia), though there was downregulation of some HSP genes in deoxygenation as well. *Eurypon* sp. 2 and *H. stellifera* both upregulated Hsp90 genes in hypoxia, but they upregulated different HSPs in anoxia: HSPA8 and Hsp90 for *Eurypon* sp. 2 and *H. stellifera*, respectively. It should be noted that Hsp90 upregulation in *H. stellifera* in anoxia was driven by only one replicate. However, in the pairwise comparison with the highest replication in this study, hypoxia vs. normoxia in *Eurypon* sp. 2, Hsp90 genes were significantly upregulated in hypoxia. Hsp90 expression was also higher in anoxia than normoxia for *Eurypon* sp. 2, but this was not significant. Based on these trends, it is possible that increased replication would result in a clearer upregulated signal from Hsp90 genes, since false negatives in gene expression analyses decrease with more replication (Schurch et al. 2016). *Eurypon* sp. 2 also upregulated HSPA8 and Hsp70 in hypoxia but downregulated Hsp40. In long term hypoxia, *T. wilhelma* exhibited a similar HSP expression pattern, upregulating Hsp90 but downregulating Hsp70 (Mills et al. 2018). Sponges under heat stress also upregulate Hsp70 (López-Legentil et al. 2008; Guzman and Conaco 2016; Webster et al. 2013) and HSP90 (Guzman and Conaco 2016), as observed under deoxygenation stress in Lough Hyne. Pollutants such as lead and zinc can also stimulate the upregulation of HSPs in sponges (Efremova et al. 2002; Schröder et al. 2006). HSPs are therefore crucial to sponge stress response, protecting and restoring proteins that could be denatured by stressors. HSP expression patterns and other key transcriptomic responses of the sponge holobionts to deoxygenation are outlined in Figure 4.

**Figure 4.**
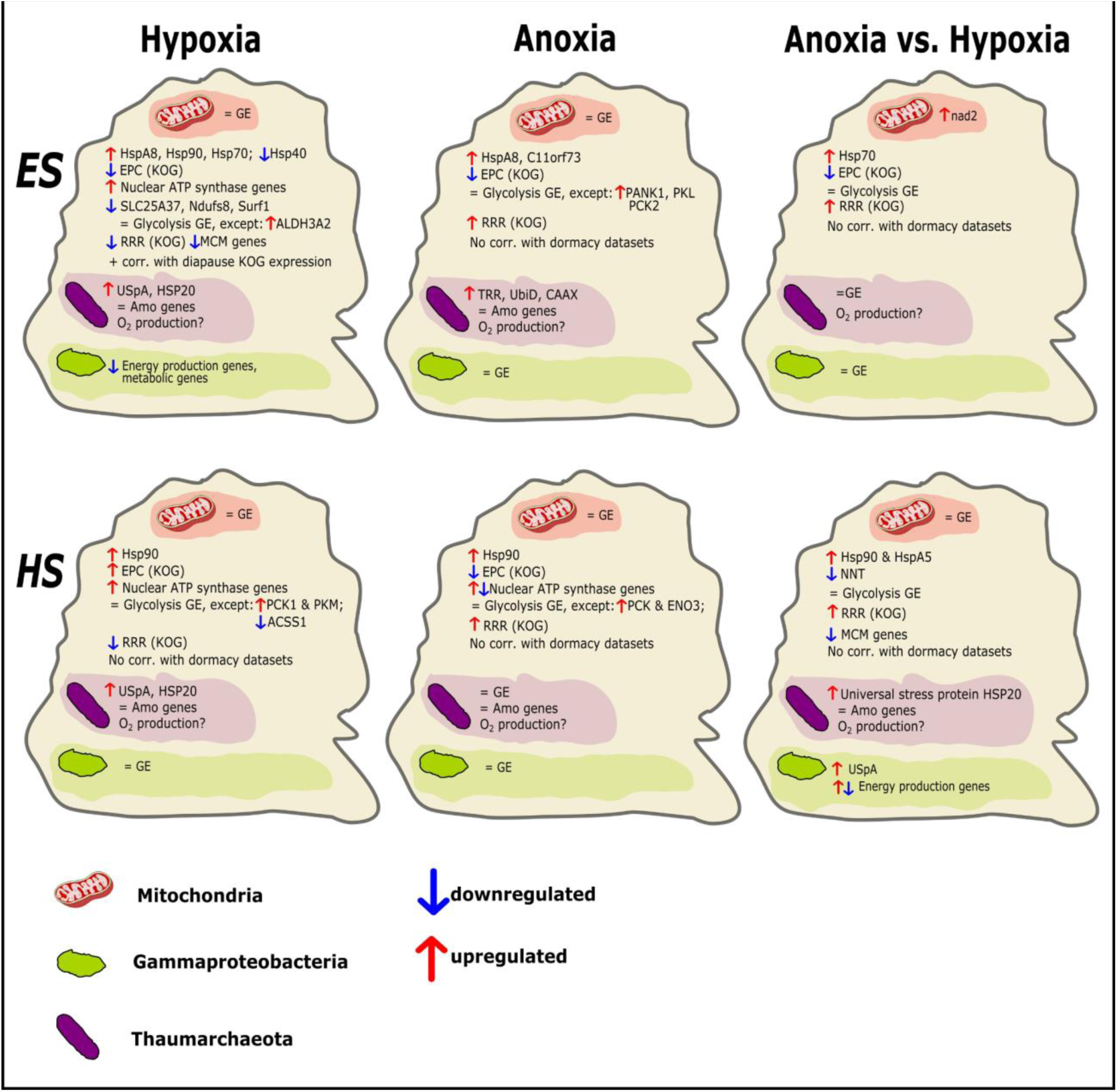
Summary of sponge holobiont responses to deoxygenation presented in a tabular format with species on the left (*Es: Eurypon* sp. 2, *Hs: H. stellifera*) and oxygen condition on the top (hypoxia, anoxia, and hypoxia vs. anoxia). The major components of the holobiont are illustrated as follows: sponge cell (tan), mitochondria (red), *Gammaproteobacteria* (green), and *Thaumarchaeota* (purple). Upregulated and downregulated genes and KOGs are indicated with red and blue arrows, respectively. Unchanged gene expression (GE) is noted by the = sign. Correlations (corr.) with external dormancy datasets are indicated along with the direction of the correlation (+ or -). The ‘energy production and conversion’ and ‘replication recombination and repair’ KOG classes are abreviated as EPC and RRR, respectively.

In metazoans with a complete HIF pathway, Hsp90 also serves to stabilize HIFa and therefore plays an important role in hypoxia tolerance (Semenza 2007). In the absence of HIFa in sponges, Hsp90 may play a greater role in oxygen homeostasis given that it is upregulated under hypoxia in all sponges that have been examined transcriptomically to date, i.e. *Eurypon* sp. 2, *H. stellifera* and *T. wilhelma,* and that Hsp90 is upregulated in *H. stellifera* under anoxia. Hsp90 could stabilize currently unknown proteins that are involved in sponge oxygen regulation or modulate transcription factors other than HIFa. Further research is needed to determine the exact role of Hsp90 in sponge oxygen homeostasis, but it presents a good candidate for a gene underlying sponge tolerance to deoxygenation.

Patterns of species and oxygen-condition-dependent expression of GO terms were observed in sponges in response to deoxygenation. GO terms such as ‘proteasome binding’ and ‘Hsp70 binding’ were upregulated in all cases of decreased oxygen in *Eurypon* sp. 2, suggesting that the species-specific response to deoxygenation in *Eurypon* sp. 2 may center around repairing or degrading damaged proteins. In *H. stellifera*, on the other hand, upregulated GO terms related to DNA repair in anoxia, including ‘damaged DNA binding’, ‘DNA binding,’ and ‘catalysis of DNA and RNA,’ so DNA repair may be more central to the deoxygenation tolerance of this species. Both protein and DNA repair upregulation might be a response to reactive oxygen species (ROS), which can become more abundant under hypoxia due to imbalances in the mitochondrial electron transport chain (Görlach et al. 2015). If there is any oxygen present in the sponges under environmental anoxia, e.g. oxygen produced by *Thaumarchaeota* (see Kraft et al. 2022, *<u>Thaumarchaeota</u>* <u>gene expression</u> below), then ROSs could cause cellular stress in anoxia. Otherwise, the presence of sulfide in the water under anoxia (Schuster et al. 2021) or ROSs other than superoxide (O_2_ ^-^) could damage proteins and DNA in anoxic conditions. HSP genes could then serve to protect and/or restore proteins damaged by sulfide or reactive oxygen species under hypoxia or anoxia.

The energy production and conversion KOG class was significantly downregulated in all oxygen conditions in both sponges, indicating potential decreases in energy production, except for hypoxia and hypoxia vs. anoxia in *H. stellifera*, when it was significantly upregulated and unchanged, respectively (Figure 1, 4). Within the energy production and conversion KOG class, however, there were species-specific responses to anoxia and hypoxia at the level of individual differentially expressed genes. Most nuclear ATP synthase genes were upregulated in hypoxia in *Eurypon* sp. 2 but were unchanged under anoxia. Similarly, two ATP synthase subunits (ATP5A1 and ATP6V0C) were upregulated in hypoxia in *H. stellifera*. In anoxia, ATP5A1 was also upregulated, but ATP6V0C was downregulated. Additionally, downregulation of SLC25A27 in *Eurypon* sp. 2 in hypoxia may increase the coupling across the mitochondrial membrane to improve ATP synthesis. We note that the electron transport chain genes Ndufs8 and Surf1 were also downregulated in hypoxia *Eurypon* sp. 2. The NNT gene, also involved in electron transport, was downregulated in anoxia vs. hypoxia in *H. stellifera*. Nevertheless, most of the genes associated with the respiration electron transport chain were not differentially expressed, and electron transport likely continued under deoxygenation. Sponges may therefore have increased ATP production under hypoxia but not under anoxia. This increase in ATP production could improve metabolic efficiency under hypoxia. In a similar way, mollusks may upregulate mitochondrial electron transport system activity and ATP synthesis capacity to improve respiration efficiency under hypoxia (Hochachka and Lutz 2001).

Glycolysis, which can produce ATP and NADH without oxygen (Stryer 1995), was expected to increase under anoxia to provide energy. However, relatively few genes directly or indirectly involved in glycolysis were differentially expressed in our sponges under deoxygenation. In *Eurypon* sp. 2, pantothenate kinase 1 (PANK1), ATP-dependent 6-phosphofructokinase (PFKL) and Phosphoenolpyruvate carboxykinase 2 (PCK2) were significantly upregulated in anoxia, and only aldehyde dehydrogenase 3 family, member a2 (ALDH3A2) was significantly upregulated in hypoxia (see Supplemental Reports, Supplemental Figure 5). In *H. stellifera*, acetyl-coenzyme A synthetase 1 (ACSS1) was significantly downregulated in hypoxia, but PCK1 and pyruvate kinase (PKM) were upregulated in hypoxia. In anoxia for *H. stellifera*, beta-enolase (ENO3) and PCK1 were upregulated. Based on these results, it is unlikely that glycolysis rates increased under anoxia and hypoxia in either species. It is more probable that glycolysis continued under anoxia and hypoxia at normal rates in both species, since gene expression in the other ∼70 genes associated with glycolysis was not significantly changed under deoxygenation (Figure 4). Either way, the sponges would have at least some access to anaerobically generated ATP through glycolysis during environmental deoxygenation.

In the KOG class ‘replication, recombination and repair’, there was significant upregulation in anoxia and anoxia vs. hypoxia, but downregulation in hypoxia (vs. normoxia) for both sponge species. In *Eurypon* sp. 2, many MCM genes involved in the initiation of DNA replication were downregulated in hypoxia but remained unchanged in anoxia. Certain aspects of DNA replication, e.g. the DNA polymerase epsilon catalytic subunit, may even be upregulated in anoxia. The pattern was less clear in *H. stellifera*, but MCM genes were downregulated in anoxia compared to hypoxia (Figure 4). These data indicated that DNA replication may decrease under hypoxia but remain stable during anoxia. In hypoxia, decreased DNA replication and, by extension, growth may be limited by increased oxidative stress and/ or increased metabolic rates under hypoxia compared to normoxia (see above).

Between the two study species, there was only one significant, positive correlation of KOG class expression between hypoxia in *Eurypon* sp. 2 and hypoxia vs. anoxia in *H. stellifera*, demonstrating the general species-specific responses of most KOGs to deoxygenation. Within a single sponge species, no correlations were seen between KOG enrichments in anoxia or hypoxia (vs. normoxia), highlighting the transcriptomic differences between responses to hypoxia and anoxia. KOG enrichments in hypoxia *versus* anoxia, on the other hand, were positively correlated with those of anoxia in *Eurypon* sp. 2 and both anoxia and hypoxia in *H. stellifera*.

When quantitatively compared to transcriptomic datasets from the sponge *T. wilhelma* using KOGMWU, transcriptomic responses in Lough Hyne sponges were only positively correlated between *T. wilhelma* under short-term anoxia and *Eurypon* sp. 2 in hypoxia vs. anoxia (Figure 1H). This positive correlation could indicate similar transcription patterns demonstrated by *T. wilhelma* and *Eurypon* sp. 2 transitioning from hypoxia to anoxia. However, no further similarities between the responses of *T. wilhelma* and Lough Hyne sponges to deoxygenation were observed. In fact, the long-term exposure of *T. wilhelma* to hypoxia was negatively correlated to KOG enrichments for both anoxia vs. hypoxia and normoxia vs. anoxia in *Eurypon* sp. 2 (Figure 1I, J), meaning that KOG expression was generally opposite between these two species for these conditions in terms of up and down regulation. The physiological responses of *T. wilhelma* and *Eurypon* sp. 2 to deoxygenation are presumably very different. These results demonstrate that large-scale transcriptomic responses to deoxygenation depend on both the studied species and oxygen concentration.

Based on KOG correlation analyses, the hypothesis that sponges are functionally dormant, i.e. temporarily ceasing growth, development, and physical activity, under anoxia was not supported (Figure 4). Sponge gene expression under anoxia did not correlate to either diapause or a dauer state in *C. elegans* (Sinha et al. 2012) or *S. mosellana* (Gong et al. 2013), respectively. Although these dormancy datasets come from distantly related invertebrates, Strader et al. (2016) demonstrated a correlation between the KOG expression patterns of a specific phenotype of coral larvae and diapause in *S. mosellana*, hypothesizing that this potential functional analog of dormancy could serve to increase larval dispersal. Therefore, the sponges of Lough Hyne could only be considered dormant under anoxia if their transcriptional method of dormancy was distinct from that of an insect, nematode and coral, which is possible but not parsimonious given that animals are monophyletic (Feuda et al. 2017). Nevertheless, it was inconclusive whether or not sponges are pumping water or feeding during anoxia, but sponges were definitively pumping under hypoxia (Schuster et al. 2021). Sponge pumping and filtering activity remain to be quantified under anoxia and hypoxia in situ.

Paradoxically, *Eurypon* sp. 2 responses to hypoxia were positively correlated to diapause in *S. mosellana* (Figure 1F, 5). Therefore, the hypoxic response in *Eurypon* sp. 2 species, rather than the anoxic response, was similar to diapause in terms of global KOG expression patterns. The replication recombination and repair KOG was also downregulated in *Eurypon* sp. 2 under hypoxia as well as in *S. mosellana* under diapause, which may partially explain the correlation between these datasets. Despite the correlation, energy production and conversion KOG expression patterns between the two species were opposing. The energy production and conversion KOG was significantly downregulated in hypoxia in *Eurypon* sp. 2 and upregulated in diapause in *S. mosselana*. These KOG enrichments are calculated from thousands of genes, so any individual gene within a KOG class might still be upregulated even if the KOG class is significantly downregulated. Therefore, it is possible that ATP production increased in hypoxia based on the upregulation of ATP synthase genes, in spite of the downregulation of the energy production and conversion KOG. Diapause can be associated with the reduction of cell division (MacRae 2010), which may occur in sponges under hypoxia but not anoxia based on MCM gene expression. In comparison, KOG enrichments under anoxia were not positively correlated to any external datasets for either sponge species (Figure 1), indicating that sponge transcriptomic responses to anoxia are relatively novel compared to those of other taxa in response to various factors in the existing literature (Dixon et al. 2015; Strader et al. 2016).

### Sponge mitochondrial gene expression during deoxygenation

A high degree of transcriptional stability under deoxygenation was observed in sponge mitochondria. Only one of the 14 mitochondrial protein-coding genes, nad2, was significantly differentially expressed in the different oxygen conditions, where it was upregulated in anoxia vs. hypoxia in *Eurypon* sp 2 (Figure 4). Both of the sponge species had higher expression of nad2 in anoxia than in normoxia or hypoxia, but this difference was only significant in *Eurypon* sp. 2.

Surprisingly little is known about differential gene expression in mitochondria, even within humans. In the limited studies available, mitochondrial gene expression was significantly different between human cancer cells and normal cells (Reznik et al. 2017) as well as between different populations of healthy pancreas tissue (Medini et al. 2021). In these studies, nad2 was significantly downregulated in most studied cancers, though it was still significantly upregulated in a few cancer types, and nad2 was also significantly differentially expressed between pancreas cell populations. Based on the general upregulation of nuclear genes associated with respiration in hypoxia, mitochondrial gene upregulation was expected in hypoxia when compared to normoxia and anoxia, but the opposite was true for nad2 in *Eurypon* sp. 2 in the current study. The other six nad genes, which code for parts of the same membrane respiratory chain complex (complex 1, NADH dehydrogenase), however, were not significantly differentially expressed in either species. In both human studies, when one mitochondrial gene was significantly differentially expressed, most other mitochondrial genes were significantly differentially expressed in the same direction (Reznik et al. 2017; Medini et al. 2021). The physiological meaning (if any) of this single, significantly differentially expressed mitochondrial gene in sponges remains to be determined. However, it is most likely that the functionality of the mitochondrial was unchanged by either anoxia or hypoxia based on gene expression results.

Compared to that of humans, even less is known about mitochondrial gene expression in marine invertebrates. Mitochondrial gene expression is usually excluded from differential expression analyses due to the differences in library preparation potentially needed for sequencing mitochondrial RNA and/or the exclusion of mitochondrial RNA from reference transcriptomes (*sensu* Meyer 2016). This dearth is partly due to the nature of mitochondrial transcription, which generates a single continuous transcript. Following transcription, however, mitochondrial transcripts are broken up into individual coding mRNA, tRNA or rRNA (D’Souza and Minczuk 2018). After the breakup, the levels of mRNAs from specific genes are controlled post-transcriptionally (D’Souza and Minczuk 2018). It is therefore possible to have different counts between the different mitochondrial genes despite their single transcript origin (Supplemental Figure 6). The consistent ‘expression’ of transcripts across oxygen conditions for each mitochondrial protein-coding gene following continuous transcription are likely to be necessary for the maintenance of mitochondrial function.

It remains unclear how sponge mitochondria maintain this transcription from an energetic standpoint under anoxia. Within the mitochondrial electron transport chain, oxygen is used as the terminal electron acceptor and is crucial for the generation of ATP through oxidative phosphorylation. Theoretically, anoxia should cause electron transport and oxidative phosphorylation in the mitochondria to cease (Chandel and Schumacker 2000). If the mitochondria are transcribing solely to maintain oxidative phosphorylation machinery until oxygen returns, then they would need to use at least enough energy (∼2 ATP per base) to transcribe for months at a time without oxygen (Lynch and Marinov 2015), and this energetic requirement would apply to the transcription of sponge nuclear genes as well. Glycolysis may provide some anaerobic ATP production, even if it is not upregulated in anoxia or hypoxia. Alternatively, to maintain ATP production through aerobic respiration, the sponges of Lough Hyne might have access to oxygen produced by their *Thaumarchaeota* symbionts, a topic explored in more detail below.

### Microbiome gene expression during deoxygenation

#### Thaumarchaeota gene expression

*Thaumarchaeota* symbionts from both sponge species exhibited significant differential gene expression under hypoxia. Only the *Thaumarchaeota* from *Eurypon* sp. 2 (OTU1) exhibited differential gene expression under anoxia, and only those of *H. stellifera* (OTU3) differentially expressed genes under anoxia vs. hypoxia. Though there were some species-specific responses, overall gene expression in both *Thaumarchaeota* was similar, supporting the hypothesis from Schuster et al. (2021) that these taxonomically similar symbionts between two sponge species might have similar functional roles.

Relatively few genes involved in anaerobic metabolism were expressed in *Thaumarchaeota* symbionts under any oxygen condition. Lactate/malate dehydrogenase, which could be involved in fermentation (Bayer et al. 2020), was upregulated in *Eurypon* sp. 2 *Thaumarchaeota*, but only in hypoxia. Genes for the other anaerobic pathways suggested by Bayer et al. 2020, i.e. succinate dehydrogenase/ fumarate reductase, nitrite reductase and sulfite reductase, were present in both *Thaumarchaeota* genomes in the present study but were not significantly differentially expressed under different oxygen conditions. The genes for nitrite reductase exhibited low expression across all samples (0–20 counts) in both *Thaumarcheaota,* and so did the dehydrogenase/ fumarate reductase gene in the *Thaumarcheaota* of *H. stellifera*. Therefore, these genes with low expression may not be used under the conditions of this study. Only three genes in the *Thaumarchaeota* of *Eurypon* sp. 2 under anoxia were significantly differentially expressed, and none of these were involved in anaerobic metabolism. No genes were differentially expressed in *H. stellifera Thaumarchaeota* under anoxia. Therefore, there was no transcriptomic evidence for the upregulation of anaerobic metabolism within *Thaumarchaeota* in anoxia.

Transcription of ammonium monooxygenase (AMO) genes did not significantly change under anoxia or hypoxia, even though the ammonia oxidation pathway is oxygen dependent. Like the sponge mitochondria, the *Thaumarchaeota* are therefore either transcribing AMO genes to be ready when oxygen returns, or they are obtaining trace oxygen for this pathway by synthesizing oxygen themselves (Kraft et al. 2022). If sponge *Thaumarchaeota* symbionts are also able to produce oxygen under anoxic conditions like their relative *N. maritimus*, then they should express AMO genes under anoxia as well as normoxia. The proposed pathway for oxygen production in *N. maritimus* involves the reduction of nitrite to nitric oxide that is dismutated into oxygen and nitrous oxide (Kraft et al. 2022). Water sampled for nitrite concentrations in Lough Hyne using established methods (Grasshoff et al. 1999) revealed that nitrite concentrations under anoxia ranged from 0.2–1.2 μM (Supplemental Table 3), increasing with depth from where seasonal anoxia and hypoxia occur. Therefore, the conditions in Lough Hyne would theoretically allow for *Thaumarchaeota* to produce oxygen under anoxia.

AsnC family proteins and nitroreductase expression was upregulated during oxygen production in *N. maritimus* (Kraft et al. 2021). In *Archaea*, AsnC proteins can regulate metabolic processes, including the switch between aerobic and anaerobic metabolisms (Lemmens et al. 2019).

Similarly, nitroreductases can have a wide range of metabolic functions (Akiva et al. 2017). *Thaumarchaeota* within *H. stellifera* upregulated a Lrp/AsnC ligand binding domain under hypoxia and in anoxia vs. hypoxia. Another AsnC type protein was also upregulated under hypoxia, and a nitroreductase was upregulated under anoxia vs. hypoxia. However, these proteins did not directly correspond to those upregulated in *N. maritimus* during oxygen production (Supplemental Figure 9), and the latter proteins are not definitively known to be involved in oxygen production, since the pathway remains unknown.

In hypoxia, both *Thaumarchaeota* upregulated the stress response genes USpA and a protein from the HSP20 family (Figure 4). It was surprising that *Thaumarchaeota* may be stressed by hypoxia since free-living thaumarchaeal Nitrosopumilales can be found in high abundances in oxygen-minimum zones (OMZs) where oxygen is either at low concentrations or below detection (Löscher et al. 2012), though some *Thaumarchaeota* OTUs decrease in abundance in suboxic and anoxic zones (Sollai et al. 2019). Members of the genus *Nitrosopumilus*, however, increase their absolute abundance in suboxic regions, but then decrease in abundance with depth in the anoxic zone (Sollai et al. 2019), indicating a preference for suboxic environments. If suboxia is also the preferred niche of sponge *Thaumarchaeota* symbionts, the upregulation of stress response genes in response to hypoxia may correspond to a rise in intracellular oxidative stress following an increase in metabolic activity in hypoxia, like their sponge hosts. In this case, these stress response proteins serve a protective and/or restorative function for other crucial proteins and structures damaged by oxidative stress.

#### Gammaproteobacteria gene expression

Gene expression in the *Gammaproteobacteria* was species and oxygen-condition dependent. In *Eurypon* sp. 2, *Gammaproteobacteria* downregulated genes in hypoxia that were associated with respiration and energy production, including cytochrome c and associated genes (Welchen et al. 2016). The *Gammaproteobacteria* also downregulated key metabolic genes in hypoxia such as glutamine synthetase and structural genes like YbiS YcfS YnhG family protein, which is involved in cell wall production (Magnet et al. 2008). So unlike sponges or *Thaumarchaeota*, *Gammaproteobacteria* in *Eurypon* sp. 2 (OTU2) may decrease metabolic activity under hypoxia, though this is likely not the case in *H. stellifera* symbionts (OTU7).

*Gammaproteobacteria* in *H. stellifera* downregulated hundreds of genes in anoxia vs. hypoxia, including USpA, so, like other members of the holobiont, hypoxia requires a stronger transcriptomic response and, therefore, may be more stressful than anoxia for these *Gammaproteobacterial* symbionts. The few upregulated genes in anoxia vs. hypoxia included genes for energy production, e.g. a transmembrane proton pump, succinyl-CoA synthetase (part of the citric acid cycle), and formate dehydrogenase (alpha subunit), although other genes associated with energy production were downregulated. Formate dehydrogenase can function in both aerobic respiration and in anaerobic respiration (Jormakka, et al. 2003), but this enzyme was not significantly upregulated in anoxia compared to normoxia or in the

*Gammaproteobacteria* of *Eurypon* sp. 2 under any condition. So, it is unlikely that this anaerobic pathway is active in sponge-associated *Gammaproteobacteria* under anoxia.

*Gammaproteobacteria* symbionts were previously hypothesized to oxidize sulfide under anoxia (Schuster et al. 2021). Although they are not present in OTU2, OTU7 from *H. stellifera* has the sulfur oxidation genes soxA, soxY, and soxZ, which could oxidize sulfide, as well as the nitrate reductase gene NAP. However, the sox genes were not expressed in any samples in the current study, and NAP was expressed at very low levels across all samples (2-28 counts). Moreover, neither symbiont exhibited significant differential gene expression in anoxia versus normoxia. Consequently, there was very little transcriptomic evidence that the anaerobic pathways in these *Gammaproteobacteria* were activated during environmental anoxia.

### Summary of potential adaptations of sponge holobionts to low oxygen

Metabolically, sponge transcription under anoxia was surprisingly similar to normoxia throughout the holobiont members, and there was no evidence of dormancy within the holobionts in anoxia (Figure 4). No clear anaerobic respiration responses were observed in the transcriptomes of sponges, their mitochondria or their two microbial symbionts under environmental anoxia. The limited significant differential expression of glycolysis genes could indicate that this metabolic process continues to comparable degrees under normoxia, hypoxia, and anoxia. Therefore, in the sponges and their microbes, glycolysis could potentially provide ATP without oxygen (Figure 4).

Hypoxia, rather than anoxia, elicited upregulation in energy production machinery such as ATP synthase (Figure 4). These potential metabolic increases suggest that within the hypoxic limits studied here (1.30-3.56 mg L^-1^), sponges perform optimally. Indeed, these limits are above those at which mortality or substantial physiological changes have been observed for sponges (Micaroni et al. 2021; Kumala et al. 2021; Mills et al. 2014). This increased energy production could be stressful, however, and the diversion of resources could limit DNA replication cell division in hypoxia (Figure 4). Somewhere between the lower limit of hypoxia (1.37 mg L^-1^) and the full anoxia studied here, sponges may no longer attempt to increase energy efficiency, and therefore elevated transcription of ATP synthase genes was not observed in anoxia.

For both the sponges and their *Thaumarcheaota* symbionts, cellular stress responses were evident under deoxygenation. Both sponge species upregulated HSP genes under anoxia and hypoxia. These genes may be crucial for sponge deoxygenation tolerance in these species (Figure 4). We hypothesize that Hsp90 plays a role in sponge oxygen homeostasis, but this needs to be tested. The *Thaumarchaeota* symbionts upregulated HSP20 and other stress response genes in anoxia and hypoxia for *H. stellifera* and hypoxia within *Eurypon* sp. 2 (Figure 4). In hypoxia, archaeal and host upregulation of stress response genes may have been a response to oxidative stress following increased metabolic activities. In anoxia, cellular stress response might result from the presence of sulfide. Additionally, post-transcriptional, metabolic changes may play a role in deoxygenation tolerance, and proteomic and metabolomic changes should be examined in future studies.

Given the largely unchanged metabolic gene expression patterns under hypoxia or anoxia in these sponges, the production of oxygen by *Thaumarchaeota* could also be crucial to holobiont survival (noted with a question mark in Figure 4). Although any oxygen produced would be at extremely low concentrations, it would have a short path of diffusion to the sponge cells, as *Thaumarchaeota* symbionts maintain high cell densities within sponges (Moeller et al. 2019). In addition, mitochondria in general can utilize very low concentrations of oxygen (500 nM) without changing their oxygen consumption rate, and cells can still respire below that threshold albeit at a lower rate (Chance 1965; Longmuir 1957; Wilson et al. 1979). In culture, oxygen generated by *Thaumarchaeota* reached 100-200 nM (Kraft et al. 2022). If sponge *Thaumarchaeota* produce oxygen at similar levels under anoxia, cellular respiration could continue in sponges under anoxia, though likely at lower respiration rates when compared to normoxia. However, given that the genes responsible are still unknown, the production of oxygen by sponge *Thaumarchaeota* needs to be verified experimentally.

### Relevance to future ocean deoxygenation

Sponges are predicted to be the ‘winners’ over other benthic organisms in the future due to their tolerance of ocean warming and acidification associated with climate change (Bell et al. 2018). Indeed, sponges have already increased in abundance in some locations while corals have decreased due to altered carbon and nutrient cycling (Pawlik et al. 2016). The additional stressor of deoxygenation that comes with climate change and coastal eutrophication has been relatively overlooked compared to ocean warming and acidification. In other marine taxa, a meta-analysis demonstrated that the incorporation of deoxygenation as a factor along with warming could substantially decrease survival times of marine organisms compared to warming alone (Vaquer-Sunyer and Duarte 2011). Since sponges may be more tolerant than other taxa to changes in all three stressors of temperature, pH and oxygen, they could indeed become more abundant in a changing ocean. However, even sponges can be lethally affected by warming, acidification (Bennett et al. 2017) and deoxygenation (Micaroni et al. 2021) in scenarios that might occur with climate change, since tolerance to these stressors is species-specific. So, surviving sponge species in a warmed, acidified, and deoxygenated ocean might be abundant, but their abundance, and diversity would likely be lower than at present. The effects of all three of these stressors together need to be understood on a much wider range of organisms to accurately predict the composition of future oceans. More importantly, the causes and effects of anthropogenic climate change need to be ameliorated as soon as possible to avoid such diversity loss.

### Relevance for animal evolution

Assuming that the first metazoans were more like an adult sponge than a ctenophore, based on morphology (Nielsen 2008) and phylogeny (Feuda et al. 2017), then some species of early sponges or proto-animals could have tolerated low environmental oxygen levels and at least some periods of benthic anoxia and euxinia in the Neoproterozoic Era (∼538.8–1,000 million years ago) when early animals, including sponges, first evolved (Dohrmann and Wörheide 2017). Lough Hyne sponges are a valuable model system for understanding early animal evolution. Early sponges could have survived under low and variable oxygen levels using mechanisms like those of Lough Hyne sponges. Therefore, contrary to longstanding hypotheses, the evolution of animals probably did not require high and stable levels of environmental oxygen.

## Acknowledgements

Field work at Lough Hyne was carried out with the permission of the National Parks and Wildlife Service of Ireland. We thank Luke Harman and Allen Whittaker of University College Cork. We also thank the Nordcee lab technicians of the Department of Biology at University of Southern Denmark for constant support in the laboratory.

## Data, scripts, code, and supplementary information availability

Raw sequence data as well as genome and transcriptome assemblies are available at NCBI under BioProject number PRJNA893197 (https://www.ncbi.nlm.nih.gov/bioproject/PRJNA893197). Scripts, code, and additional data are available here: https://zenodo.org/badge/latestdoi/573040118. Supplemental figures and tables are available on BioRxiv under ‘Supplementary material.’

## Taxonomic authority citations

1 Grant, 1836

2 Pallas, 1766

3 Bowerbank, 1858

4 Dendy, 1905

5 Grant, 1826

6 Johnston, 1842

7 Kelly-Borges and Bergquist, 1997

8 Bergquist, 1968

9 Schmidt, 1870

10 Lévi, 1953

11 Topsent, 1928

12 Bosc, 1802

13 Bowerbank, 1864

