## Supplemental material for "Transcriptomic responses of sponge holobionts to in situ, seasonal anoxia and hypoxia"

**Supplemental Table 1.** Collection metadata for all sponge samples used in this study.

| Species | Individual | Depth (m) | Oxygen concentration (mg L-1) | Oxygen condition | Temperature (°C) | Season | Year | Month | Day |
|---|---|---|---|---|---|---|---|---|---|
| *Eurypon sp. 2* | DC33 | 19 | 9.51 | Normoxic | 15.64 | Summer | 2018 | 8 | 8 |
| *Eurypon sp. 2* | DC39 | 14.8 | 9.51 | Normoxic | 15 | Summer | 2018 | 8 | 8 |
| *Eurypon sp. 2* | DC85 | 24 | 11.81 | Normoxic | 11.37 | Spring | 2019 | 4 | 9 |
| *Eurypon sp. 2* | DC93 | 27 | 12.20 | Normoxic | 12.69 | Spring | 2019 | 4 | 10 |
| *Eurypon sp. 2* | DC97 | 27 | 11.70 | Normoxic | 11.55 | Spring | 2019 | 4 | 11 |
| *Eurypon sp. 2* | DC135 | 20 | 9.25 | Normoxic | 14.73 | Summer | 2019 | 7 | 26 |
| *Eurypon sp. 2* | DC136 | 20 | 9.25 | Normoxic | 14.73 | Summer | 2019 | 7 | 26 |
| *Eurypon sp. 2* | DC108 | 27 | 2.36 | Hypoxic | 13.04 | Summer | 2019 | 7 | 23 |
| *Eurypon sp. 2* | DC112 | 27 | 1.70 | Hypoxic | 13.62 | Summer | 2019 | 7 | 24 |
| *Eurypon sp. 2* | DC114 | 27 | 1.70 | Hypoxic | 13.62 | Summer | 2019 | 7 | 24 |
| *Eurypon sp. 2* | DC118 | 27 | 1.70 | Hypoxic | 13.62 | Summer | 2019 | 7 | 24 |
| *Eurypon sp. 2* | DC123 | 27 | 1.71 | Hypoxic | 13.80 | Summer | 2019 | 7 | 25 |
| *Eurypon sp. 2* | DC131 | 27 | 1.37 | Hypoxic | 13.68 | Summer | 2019 | 7 | 26 |
| *Eurypon sp. 2* | DC110 | 27 | 2.36 | Hypoxic | 13.04 | Summer | 2019 | 7 | 23 |
| *Eurypon sp. 2* | DC24 | 27 | 0.00 | Anoxic | 10.80 | Summer | 2018 | 8 | 7 |
| *Eurypon sp. 2* | DC56 | 27 | 0.00 | Anoxic | 11.88 | Summer | 2018 | 8 | 9 |
| *H. stellifera* | DC129 | 20 | 9.11 | Normoxic | 14.72 | Summer | 2019 | 7 | 25 |
| *H. stellifera* | DC105 | 26.3 | 2.36 | Hypoxic | 13.04 | Summer | 2019 | 7 | 23 |
| *H. stellifera* | DC107 | 29 | 2.36 | Hypoxic | 13.04 | Summer | 2019 | 7 | 23 |
| *H. stellifera* | DC133 | 27 | 1.37 | Hypoxic | 13.68 | Summer | 2019 | 7 | 26 |
| *H. stellifera* | DC121 | 27 | 1.71 | Hypoxic | 13.80 | Summer | 2019 | 7 | 25 |
| *H. stellifera* | DC116 | 27 | 1.70 | Hypoxic | 13.62 | Summer | 2019 | 7 | 24 |
| *H. stellifera* | DC54 | 27 | 0.00 | Anoxic | 11.88 | Summer | 2018 | 9 | 8 |
| *H. stellifera* | DC55 | 27 | 0.00 | Anoxic | 11.88 | Summer | 2018 | 9 | 8 |

**Supplemental Table 2**. Table summarizing all stats for reference transcriptomes and metagenomes.

| | ***Eurypon* sp. 2 holobiont** | | | | ***H. stellifera* holobiont** | | | |
|---|---|---|---|---|---|---|---|---|
| Sequence type | mRNA | Total RNA | DNA | | mRNA | Total RNA | DNA | |
| N raw reads (x $10^6$) | 1,120 | 1,420 | 152 | | 793 | 788 | 99.8 | |
| N qual filtered: PE, SE (x $10^6$) | 55.2, 8.29 | NA | NA | | 19.7, 3.10 | NA | NA | |
| N contigs holobiont | 147,136 | NA | 232,756 | | 128,943 | NA | 88,622 | |
| ***Target reference (bin) only:*** | **Sponge** | **Mito-chondria** | ***Thaum-archaeota*** | ***Gamma-proteo-bacteria*** | **Sponge** | **Mito-chondria** | ***Thaum-archaeota*** | ***Gamma-proteo-bacteria*** |
| N contigs target | 127,979 | 14 | 370 | 983 | 110,497 | 14 | 157 | 546 |
| Mean GC content target | 51.0 | 28.8 | 58.6 | 58.2 | 49.2 | 25.5 | 33.2 | 57.2 |
| N genes | 49,911 | 14 | 1,556 | 2,751 | 62,494 | 14 | 1,551 | 3,057 |
| Mean contig length (bp) | 1,047 | NA | 4,966 | 6,649 | 1,135 | NA | 12,232 | 12,473 |
| N50 (bp) | 1,283 | NA | 11,143 | 14,923 | 1,545 | NA | 17,952 | 28,974 |
| % Annotated | 39.7 | 100 | 69.2 | 81.5 | 37.0 | 100 | 67.7 | 84.8 |
| *BUSCOs*: | - | - | - | - | - | - | - | - |
| N complete (%) | 94.7 | NA | 90.0 | 95.2 | 95.9 | NA | 97.0 | 96.5 |
| N partial (%) | 96.8 | NA | 92.8 | 96.7 | 96.9 | NA | 97.7 | 97.0 |
| N missing (%) | 3.17 | NA | 7.20 | 3.23 | 3.07 | NA | 2.35 | 3.01 |

**Supplemental Table 3.** Nitrite concentrations with depth in Lough Hyne, sampled on 26/7/2019 when conditions at the sponge sampling depth were hypoxic. Anoxia occurred at a depth of 33 m.

| Depth (m) | 1 | 5 | 10 | 15 | 20 | 25 | 27 | 30 | 32 | 34 | 35 | 36 | 38 |
|---|---|---|---|---|---|---|---|---|---|---|---|---|---|
| Nitrite concentration (µM) | 0 | 0 | 0 | 0 | 0 | 0 | 0 | 0 | 0 | 0.31 | 0.81 | 0.76 | 1.11 |

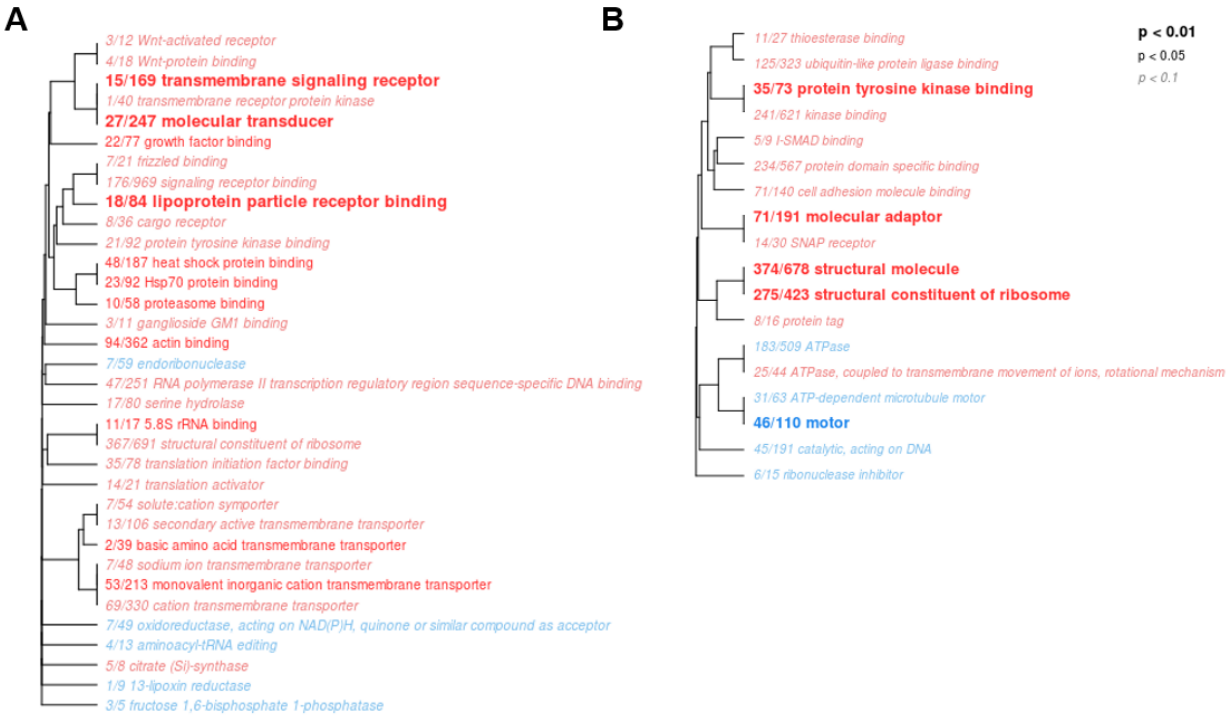


**Supplemental Figure 1.** Significantly enriched GO categories of biological processes in A. *Eurypon* sp. 2 (34 GO terms) and B. *Hymerphia stellifera* (18 GO terms) in normoxia vs. hypoxia. Red GO terms are significantly upregulated in hypoxia, and blue terms are downregulated. The font and text size correspond to the p values (see legend) of Fisher's exact tests of GO category enrichment. The numbers in front of the GO categories correspond to the number of significantly differentially expressed genes (numerator) based on treatment out of the total number of genes in the dataset belonging to that GO category (denominator). GO categories are clustered hierarchically based on shared genes between the categories, meaning that those on the same branch are subsets of one another.

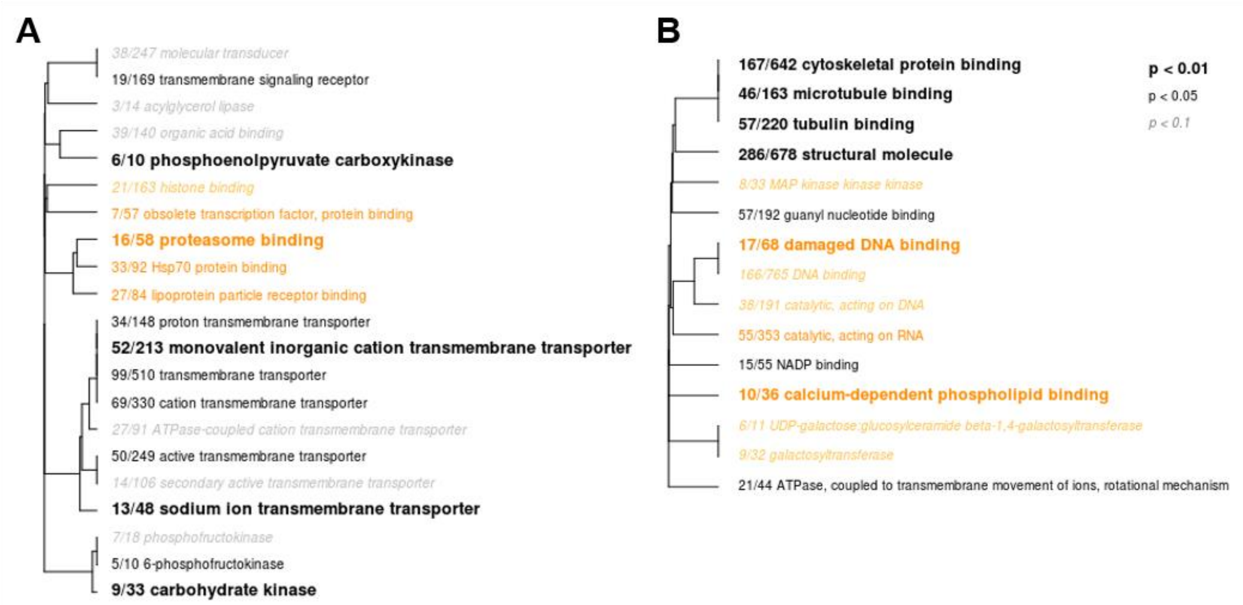


**Supplemental Figure 2.** Significantly enriched GO categories of biological processes in A. *Eurypon* sp. 2 (12 GO terms) and B. *Hymerphia stellifera* (17 GO terms) in normoxia vs. anoxia. Orange GO terms are significantly upregulated in anoxia, and black terms are downregulated. The font and text size correspond to the p values (see legend) of Fisher's exact tests of GO category enrichment. The numbers in front of the GO categories correspond to the number of significantly differentially expressed genes (numerator) based on treatment out of the total number of genes in the dataset belonging to that GO category (denominator). GO categories are clustered hierarchically based on shared genes between the categories, meaning that those on the same branch are subsets of one another.

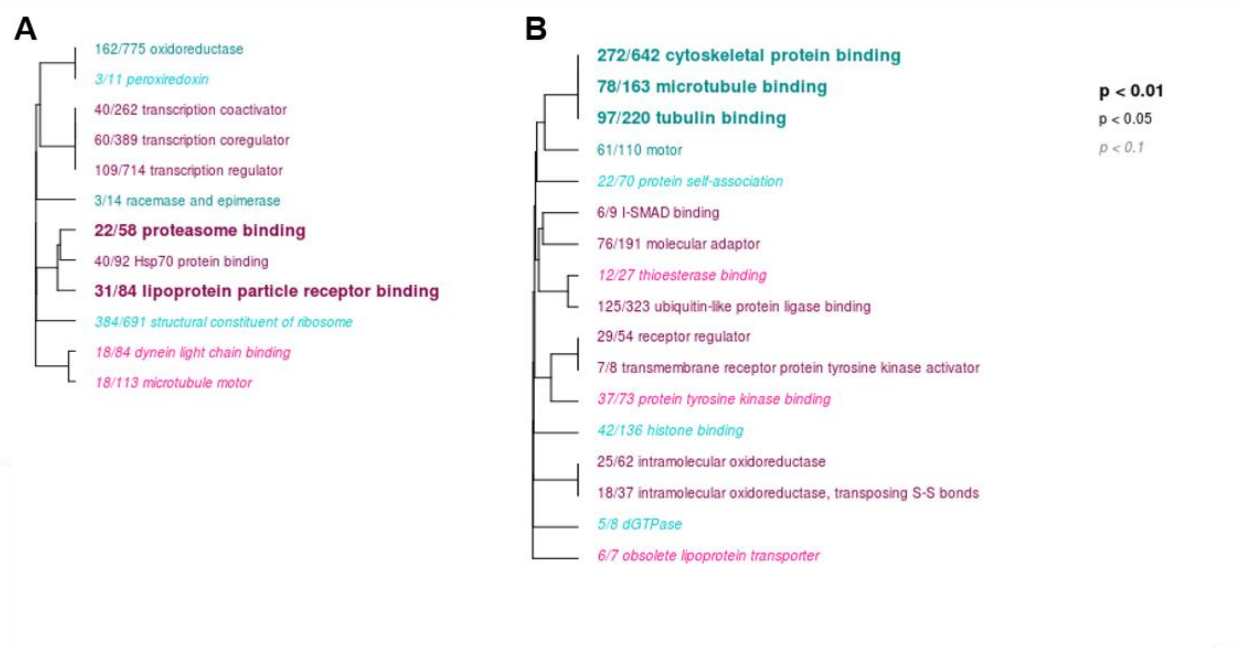


**Supplemental Figure 3.** Significantly enriched GO categories of biological processes in A. *Eurypon* sp. 2 (12 GO terms) and B. *Hymerphia stellifera* (17 GO terms) in hypoxia vs. anoxia. Pink GO terms are significantly upregulated in anoxia, and cyan terms are downregulated. The font and text size correspond to the p values (see legend) of Fisher's exact tests of GO category enrichment. The numbers in front of the GO categories correspond to the number of significantly differentially expressed genes (numerator) based on treatment out of the total number of genes in the dataset belonging to that GO category (denominator). GO categories are clustered hierarchically based on shared genes between the categories, meaning that those on the same branch are subsets of one another.

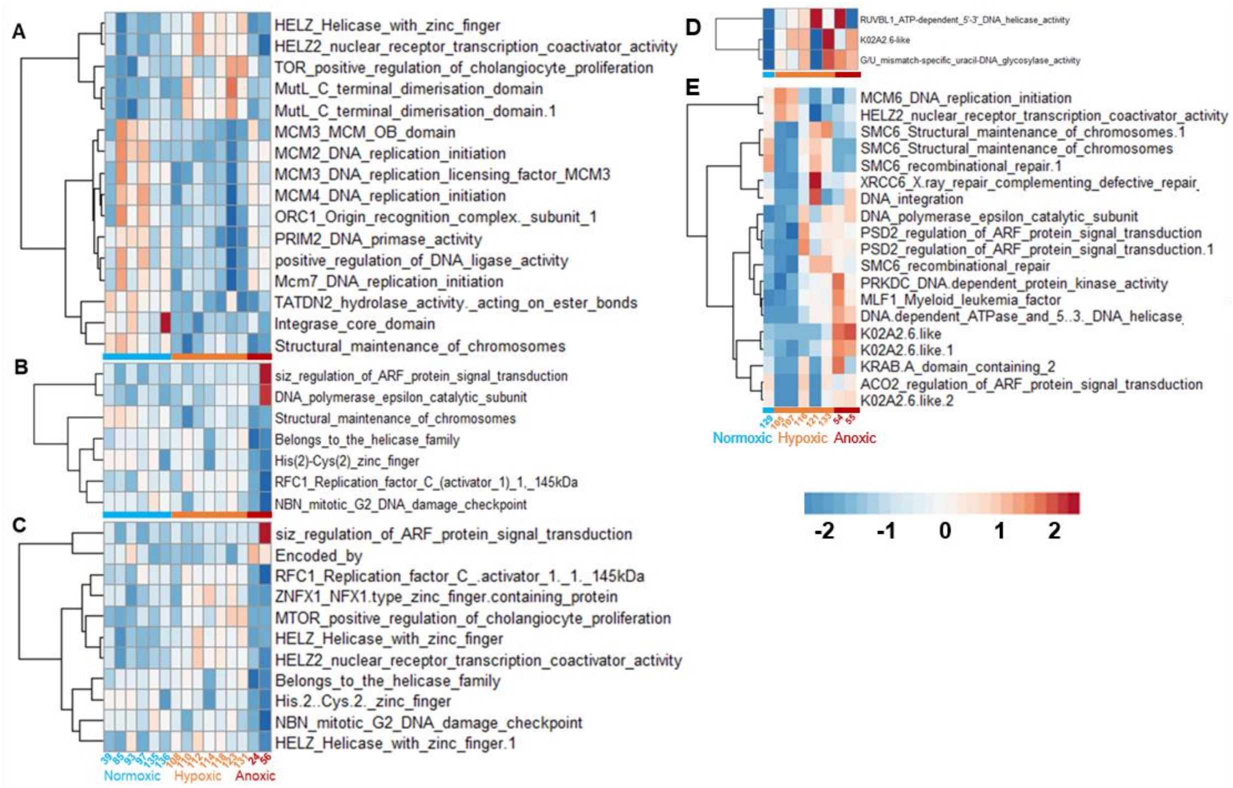


**Supplemental Figure 4.** All significantly differentially expressed genes within the KOG class 'replication, recombination and repair' (RRR) in both sponge species. For *Eurypon* sp. 2 the three pairwise comparisons are shown in A (normoxia vs. hypoxia), B (hypoxia vs. normoxia), and C (anoxia vs. hypoxia). Significantly differentially expressed RRR genes in *H. stellifera* are shown in D and E for normoxia vs. hypoxia and hypoxia vs. anoxia, respectively. Samples collected under normoxia, hypoxia and anoxia are shown in blue, orange and red, respectively. Heatmaps are scaled based on row (gene) mean, i.e. Z-scores.

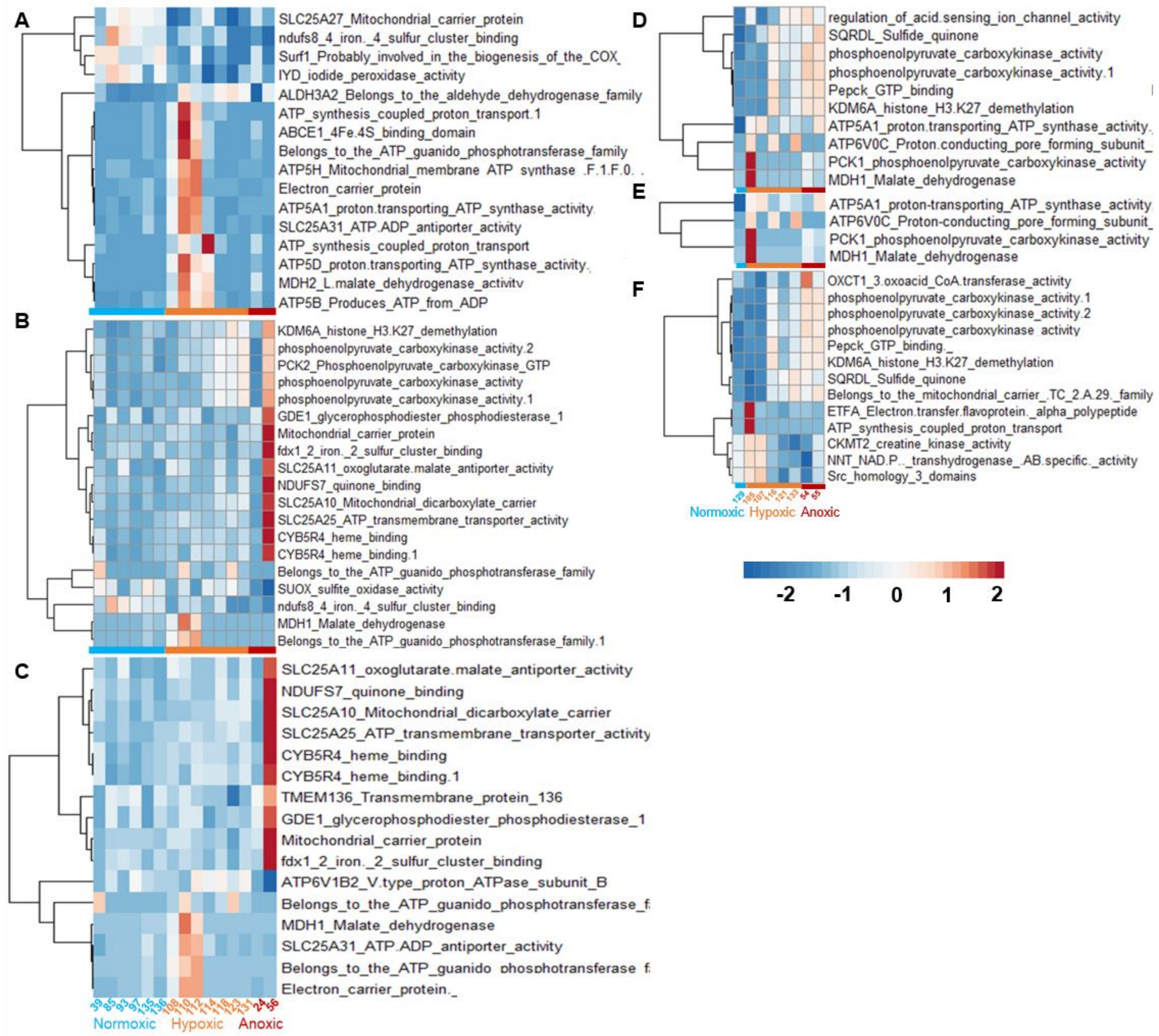


**Supplemental Figure 5.** All significantly differentially expressed genes within the KOG class 'energy production and conversion' (EPC) in both sponge species. For *Eurypon* sp. 2 the three pairwise comparisons are shown in A (normoxia vs. hypoxia), B (hypoxia vs. normoxia), and C (anoxia vs. hypoxia). Significantly differentially expressed EPC genes in *H. stellifera* are shown in D-F for normoxia vs. hypoxia, normoxia vs. anoxia, and hypoxia vs. normoxia, respectively. Samples collected under normoxia, hypoxia and anoxia are shown in blue, orange and red, respectively. Heatmaps are scaled based on row (gene) mean, i.e. Z-scores.

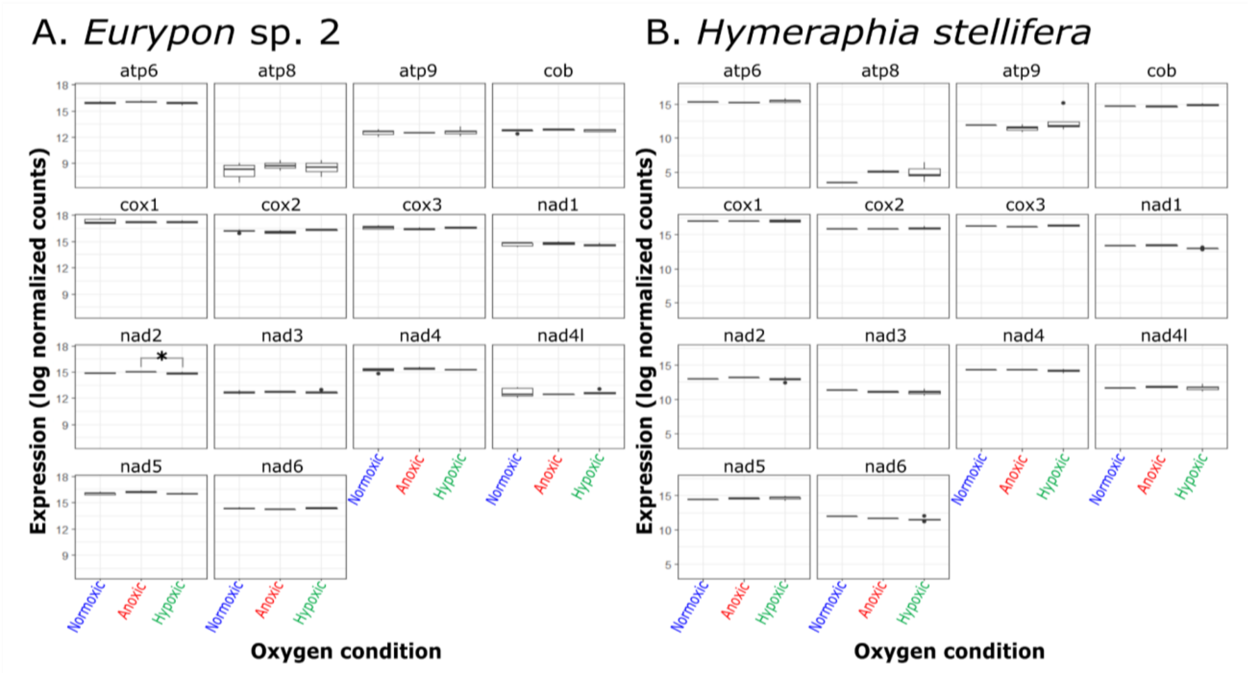


**Supplemental Figure 6**. Mitochondrial gene expression under different oxygen conditions in A. *Eurypon* sp. 2 and B. *Hymeraphia stellifera*. Color code corresponds to Schuster, Strehlow et. al. 2021. Asterisk (*) indicates a significant difference in the nad2 gene of *Eurypon* sp. 2.

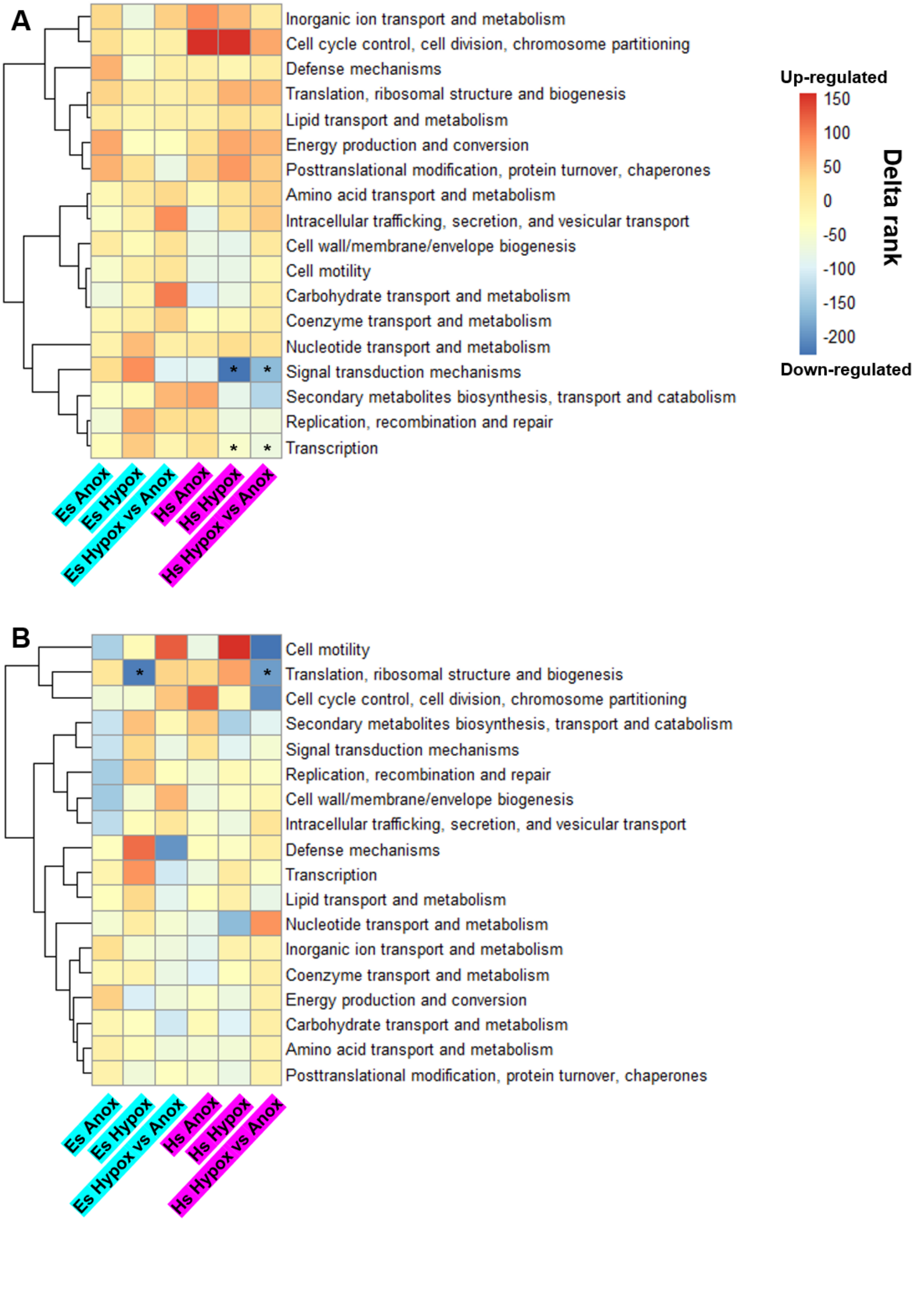
A
Inorganic ion transport and metabolism
Cell cycle control, cell division, chromosome partitioning
Defense mechanisms
Translation, ribosomal structure and biogenesis
Lipid transport and metabolism
Energy production and conversion
Posttranslational modification, protein turnover, chaperones
Amino acid transport and metabolism
Intracellular trafficking, secretion, and vesicular transport
Cell wall/membrane/envelope biogenesis
Cell motility
Carbohydrate transport and metabolism
Coenzyme transport and metabolism
Nucleotide transport and metabolism
Signal transduction mechanisms
Secondary metabolites biosynthesis, transport and catabolism
Replication, recombination and repair
Transcription
Es Anox
Es Hypox
Es Hypox vs Anox
Hs Anox
Hs Hypox
Hs Hypox vs Anox
Up-regulated
150
100
50
0
-50
-100
-150
-200
Down-regulated
Delta rank
B
Cell motility
Translation, ribosomal structure and biogenesis
Cell cycle control, cell division, chromosome partitioning
Secondary metabolites biosynthesis, transport and catabolism
Signal transduction mechanisms
Replication, recombination and repair
Cell wall/membrane/envelope biogenesis
Intracellular trafficking, secretion, and vesicular transport
Defense mechanisms
Transcription
Lipid transport and metabolism
Nucleotide transport and metabolism
Inorganic ion transport and metabolism
Coenzyme transport and metabolism
Energy production and conversion
Carbohydrate transport and metabolism
Amino acid transport and metabolism
Posttranslational modification, protein turnover, chaperones
Es Anox
Es Hypox
Es Hypox vs Anox
Hs Anox
Hs Hypox
Hs Hypox vs Anox

**Supplemental Figure 7.** Gene expression profiles of sponge associated microbes under varied oxygen concentrations compared to each other. A. Clustered heatmap (based on correlation) of COG class (rows) enrichments for upregulated or downregulated genes for *Thaumarchaeota* symbionts. B. Clustered heatmap of COG class enrichments in *Gammaproteobacteria* symbionts. Highlighted text colors correspond to the sponge species to which each symbiont is associated. (Purple: *Hymeraphia stellifera* [Hs], Cyan: *Eurypon* sp. 2 [Es]). Up or down regulation with respect to normoxia is indicated in 'Hypox' (hypoxia) and 'Anox' (anoxia) columns, and differential regulation with respect to hypoxia is shown in 'Hypox vs. Anox' columns, i.e. upregulated COGs (red delta ranks) are upregulated in anoxia with respect to hypoxia. Within each column on the heatmap, COGs that are significantly enriched (FDR-adjusted $p < 0.05$) are outlined in black boxes.

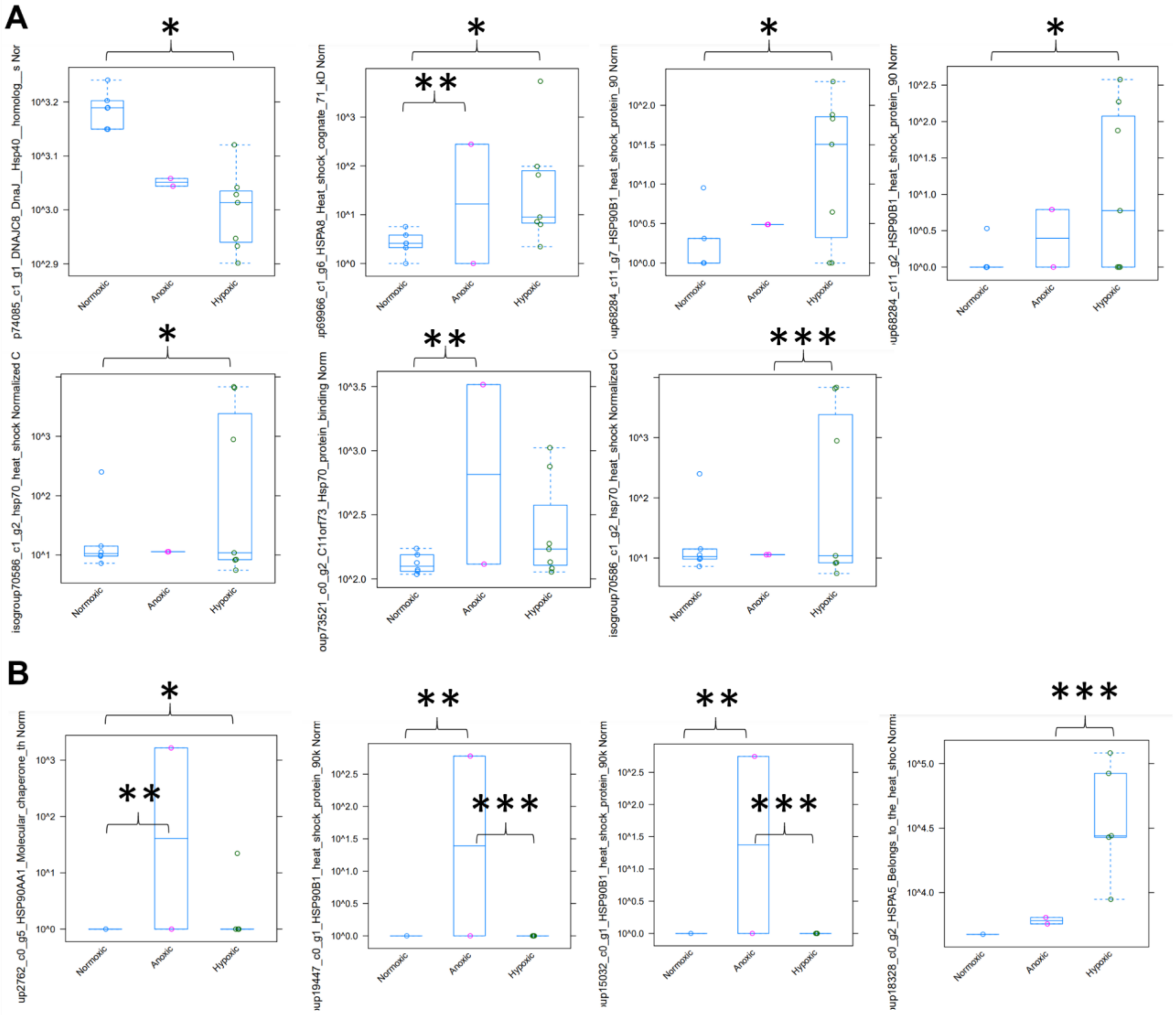


**Supplemental Figure 8.** Significantly differentially expressed heat shock protein (HSP) genes in sponges in response to deoxygenation. A. *Eurypon* sp. 2. B. *H. stellifera*. Asterisks: *, ** and

*** indicate significant differences in expression level between normoxia vs. hypoxia, normoxia vs. anoxia, and anoxia vs. hypoxia, respectively.

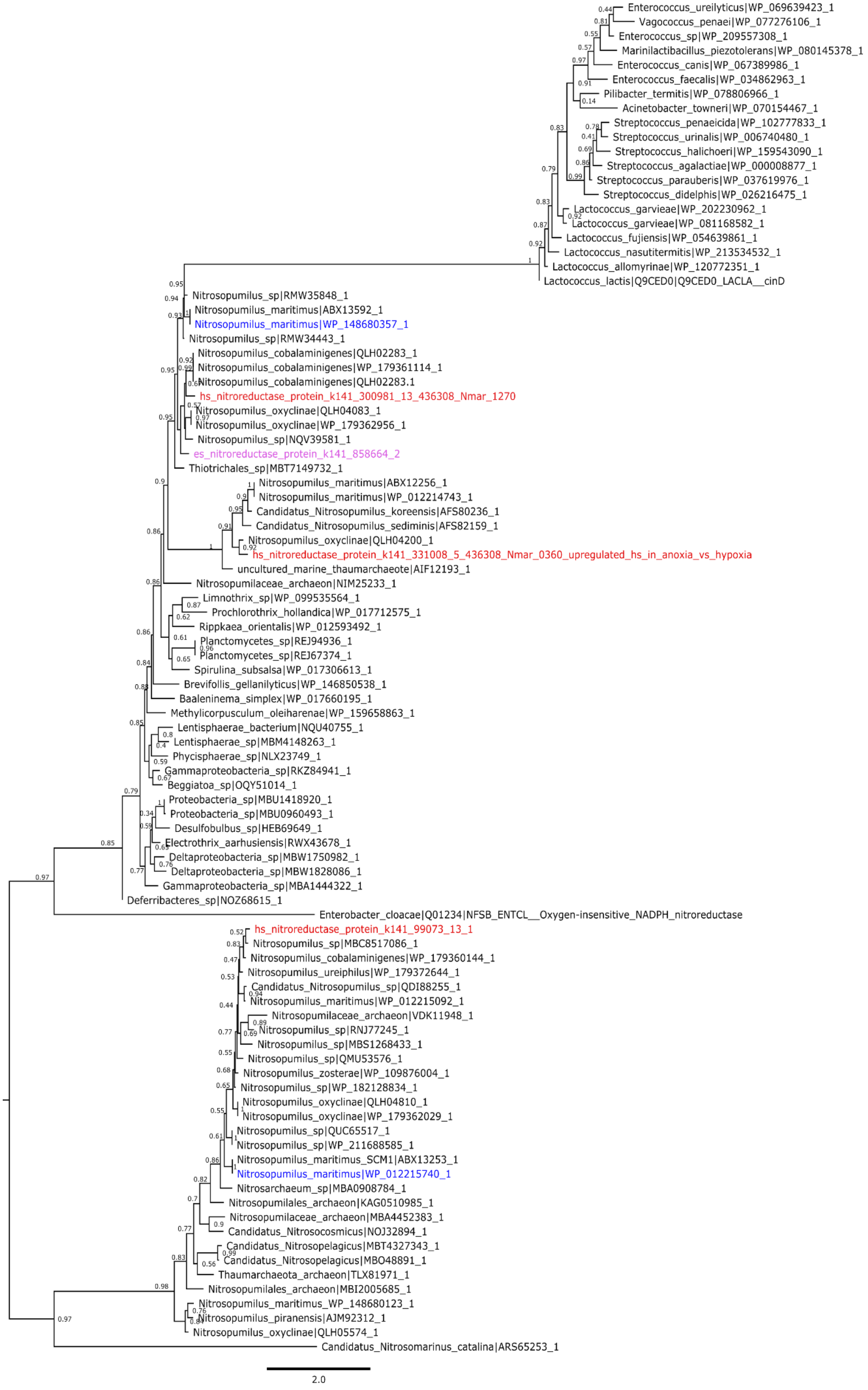

Enterococcus_ureilyticus|WP_069639423_1
Vagococcus_penaei|WP_077276106_1
Enterococcus_sp|WP_209557308_1
Marinilactibacillus_piezotolerans|WP_080145378_1
Enterococcus_canis|WP_067389986_1
Enterococcus_faecalis|WP_034862963_1
Pilibacter_termitis|WP_078806966_1
Acinetobacter_towneri|WP_070154467_1
Streptococcus_penaeicida|WP_102777833_1
Streptococcus_urinalis|WP_006740480_1
Streptococcus_halichoeri|WP_159543090_1
Streptococcus_agalactiae|WP_000008877_1
Streptococcus_parauberis|WP_037619976_1
Streptococcus_didelphis|WP_026216475_1
Lactococcus_garvieae|WP_202230962_1
Lactococcus_garvieae|WP_081168582_1
Lactococcus_fujiensis|WP_054639861_1
Lactococcus_nasutitermitis|WP_213534532_1
Lactococcus_allomyrinae|WP_120772351_1
Lactococcus_lactis|Q9CED0|Q9CED0_LACLA__cinD
Nitrosopumilus_sp|RMW35848_1
Nitrosopumilus_maritimus|ABX13592_1
Nitrosopumilus_maritimus|WP_148680357_1
Nitrosopumilus_sp|RMW34443_1
Nitrosopumilus_cobalaminigenes|QLH02283_1
Nitrosopumilus_cobalaminigenes|WP_179361114_1
Nitrosopumilus_cobalaminigenes|QLH02283.1
hs_nitroreductase_protein_k141_300981_13_436308_Nmar_1270
Nitrosopumilus_oxyclinae|QLH04083_1
Nitrosopumilus_oxyclinae|WP_179362956_1
Nitrosopumilus_sp|NQV39581_1
es_nitroreductase_protein_k141_858664_2
Thiotrichales_sp|MBT7149732_1
Nitrosopumilus_maritimus|ABX12256_1
Nitrosopumilus_maritimus|WP_012214743_1
Candidatus_Nitrosopumilus_koreensis|AFS80236_1
Candidatus_Nitrosopumilus_sediminis|AFS82159_1
Nitrosopumilus_oxyclinae|QLH04200_1
hs_nitroreductase_protein_k141_331008_5_436308_Nmar_0360_upregulated_hs_in_anoxia_vs_hypoxia
uncultured_marine_thaumarchaeote|AIF12193_1
Nitrosopumilaceae_archaeon|NIM25233_1
Limnothrix_sp|WP_099535564_1
Prochlorothrix_hollandica|WP_017712575_1
Rippkaea_orientalis|WP_012593492_1
Planctomycetes_sp|REJ94936_1
Planctomycetes_sp|REJ67374_1
Spirulina_subsalsa|WP_017306613_1
Brevifollis_gellanilyticus|WP_146850538_1
Baaleninema_simplex|WP_017660195_1
Methylicorpusculum_oleiharenae|WP_159658863_1
Lentisphaerae_bacterium|NQU40755_1
Lentisphaerae_sp|MBM4148263_1
Phycisphaerae_sp|NLX23749_1
Gammaproteobacteria_sp|RKZ84941_1
Beggiatoa_sp|OQY51014_1
Proteobacteria_sp|MBU1418920_1
Proteobacteria_sp|MBU0960493_1
Desulfobulbus_sp|HEB69649_1
Electrothrix_aarhusiensis|RWX43678_1
Deltaproteobacteria_sp|MBW1750982_1
Deltaproteobacteria_sp|MBW1828086_1
Gammaproteobacteria_sp|MBA1444322_1
Deferribacteres_sp|NOZ68615_1
Enterobacter_cloacae|Q01234|NFSB_ENTCL__Oxygen-insensitive_NADPH_nitroreductase
hs_nitroreductase_protein_k141_99073_13_1
Nitrosopumilus_sp|MBC8517086_1
Nitrosopumilus_cobalaminigenes|WP_179360144_1
Nitrosopumilus_ureiphilus|WP_179372644_1
Candidatus_Nitrosopumilus_sp|QDI88255_1
Nitrosopumilus_maritimus|WP_012215092_1
Nitrosopumilaceae_archaeon|VDK11948_1
Nitrosopumilus_sp|RNJ77245_1
Nitrosopumilus_sp|MBS1268433_1
Nitrosopumilus_sp|QMU53576_1
Nitrosopumilus_zosterae|WP_109876004_1
Nitrosopumilus_sp|WP_182128834_1
Nitrosopumilus_oxyclinae|QLH04810_1
Nitrosopumilus_oxyclinae|WP_179362029_1
Nitrosopumilus_sp|QUC65517_1
Nitrosopumilus_sp|WP_211688585_1
Nitrosopumilus_maritimus_SCM1|ABX13253_1
Nitrosopumilus_maritimus|WP_012215740_1
Nitrosarchaeum_sp|MBA0908784_1
Nitrosopumilales_archaeon|KAG0510985_1
Nitrosopumilaceae_archaeon|MBA4452383_1
Candidatus_Nitrosocosmicus|NOJ32894_1
Candidatus_Nitrosopelagicus|MBT4327343_1
Candidatus_Nitrosopelagicus|MBO48891_1
Thaumarchaeota_archaeon|TLX81971_1
Nitrosopumilales_archaeon|MBI2005685_1
Nitrosopumilus_maritimus_WP_148680123_1
Nitrosopumilus_piranensis|AJM92312_1
Nitrosopumilus_oxyclinae|QLH05574_1
Candidatus_Nitrosomarinus_catalina|ARS65253_1
2.0

**Supplemental Figure 9.** Maximum Likelihood (ML) phylogeny based on nitroreductase protein alignment. Numbers on branches indicate bootstrap support (BS) values. The position of the nitroreductase proteins from *Hymeraphia stellifera* (hs) are highlighted in red and of *Eurypon* sp. 2 (es) in pink. Sequences from Kraft et al. (2022) are highlighted in blue.
